# Classical HLA Allele and Haplotype Frequency Estimates in US Populations

**DOI:** 10.64898/2026.04.09.717537

**Authors:** Loren Gragert, Abeer Madbouly, Pradeep Bashyal, Kim Wadsworth, Jane Kempenich, Yung-Tsi Bolon, Martin Maiers

## Abstract

The human leukocyte antigen (HLA) system is the primary determinant of donor selection in allogeneic hematopoietic cell transplantation (HCT) and plays a central role in solid organ transplantation, immune-mediated disease studies, evolutionary population genetics, and immunotherapy. Large-scale sampling of registry participants reflecting major US ancestry groups allows for characterization of the complex landscape of HLA haplotype diversity for the classical HLA class I (*HLA-A, HLA-B, HLA-C*) and HLA class II (*HLA-DRB1, HLA-DRB3, HLA-DRB4, HLA-DRB5, HLA-DQA1, HLA-DQB1, HLA-DPA1, and HLA-DPB1*) genes. Here we present nine-locus classical HLA allele and haplotype frequency estimates for five broad (Black, White, Asian or Pacific Islander, Hispanic and Native American) and 21 detailed US populations based on 9,671,082 donors with targeted genotyping by DNA-based methods. Frequency estimation used an expectation–maximization (EM) framework specifically adapted to handle mixed-resolution and ambiguous HLA genotyping data. Advancements in next-generation sequencing provide extensive HLA genotyping, offering new insights into the haplotype structure and diversity of the human MHC complex, expanding knowledge especially for HLA class II haplotypes. Population analyses reveal that the most common high-resolution haplotypes are predominantly population-specific, with only three haplotypes shared across the top-100 lists of all five broad population groups, and that Black populations exhibit the greatest nine-locus haplotypic diversity, a pattern that persists after controlling for differences in registry sample size. These frequencies, derived from the largest US cohort to date, support clinical decision-making and research in histocompatibility, immunogenetics, and transplantation and are publicly available at https://zenodo.org/records/17966993.

## INTRODUCTION

The classical human leukocyte antigen (HLA) genes are the most polymorphic genes in the human genome and play an integral role in the adaptive immune response [1]. Classical HLA class I genes (*HLA-A, HLA-B, HLA-C*) are expressed on nearly all nucleated cells. They present peptide antigens to CD8+ T cells. HLA class I molecules are also ligands for killer immunoglobulin receptor (KIR) receptors on natural killer (NK) cells. Classical HLA class II genes (*HLA-DRB1, HLA-DRB3, HLA-DRB4, HLA-DRB5, HLA-DQA1, HLA-DQB1, HLA-DPA1, HLA-DPB1*) present peptide antigens to CD4+ T cells and help initiate an antibody response through linked recognition with B cells [2].

HLA genetic diversity in human populations has been shaped by forces of evolutionary selection and population history. Global populations exhibit remarkable differences in HLA allele and haplotype frequency. HLA haplotype frequency estimates have numerous applications in multiple donor registry operations like modeling bone marrow registry match likelihoods [3], focused recruitment of registry donors, statistical imputation [4-6] and donor/recipient matching [7], genetic association studies of immune-mediated diseases [8-12], T-cell epitopes for immunotherapy [13-15] and vaccine coverage [16, 17], and evolutionary population genetics [18-22].

Because HLA loci span a three-megabase region of chromosome 6 and are genotyped by targeted sequencing of HLA genes without intervening genomic sequence, haplotype phasing cannot be determined directly from the typing assay. Instead, computational maximum likelihood-based methods provide population-level haplotype frequency estimates [23, 24]. Through targeted HLA genotyping of stem cell registry volunteers, the NMDP (*formerly National Marrow Donor Program*) database provides a deep sampling of US population HLA diversity, including indigenous populations of the US. Genotyping of HLA genes has expanded in resolution and number of genes with the advent of lower cost next-generation sequencing (NGS) based methods. Historical genotyping on donors recruited in the 1990s and 2000s had high allelic and genotypic ambiguity and did not include all classical HLA genes. However, with continued evolution of transplant clinical outcomes research [13, 25-27] and donor selection guidelines [28-32], accrual of genotyping was extended for NMDP volunteer donors to include *HLA-DQA1, HLA-DPA1*, and *HLA-DPB1* to expand haplotype frequency estimation to all classical HLA genes.

Building on the work of Kollman et al. [23] and Gragert et al. [24], we extended the NMDP expectation-maximization (EM) algorithm, that simultaneously resolves both phase and allelic ambiguity, to estimate up to nine-locus haplotypes encompassing all classical HLA loci. Previously, there were some substantial computational complexities that rendered the estimation of nine-locus frequencies difficult or prohibitive as the sample sizes and HLA genetic diversity grew for US populations. Here we describe haplotype frequency estimates from a new, streamlined implementation of the EM pipeline that addresses previous computational limitations and can easily be extended to estimate beyond nine-locus haplotypes. This is timely due to increases in both the inclusion of more HLA class II genes in routine HLA genotyping and the consideration of these genes in donor selection. The presented haplotype frequencies are estimated from the largest published US-based sample to date (approximately 10 million samples). These frequencies are used as a reference dataset that benefits multiple research and clinical domains and applications such as matching for stem-cell transplant, resolving gaps and ambiguities in HLA genotyping data for stem-cell and solid organ donors and patients, donor registry planning and match rate projections, guiding optimal registry strategy, disease association studies, vaccine design and beyond.

## METHODS

### Study Population

A sample of 9,671,082 volunteer registry donors typed at various DNA methods and divided into 21 detailed and five broad populations (Tables 1 and 2) was used to estimate nine-locus haplotype frequencies for each of the studied groups. Inclusion criteria were all US donors listed in the NMDP registry that were first genotyped by PCR-DNA methods, at minimum at *HLA-A, HLA-B*, and *HLA-DRB1* genes as of July 2024.

**Table 1.**
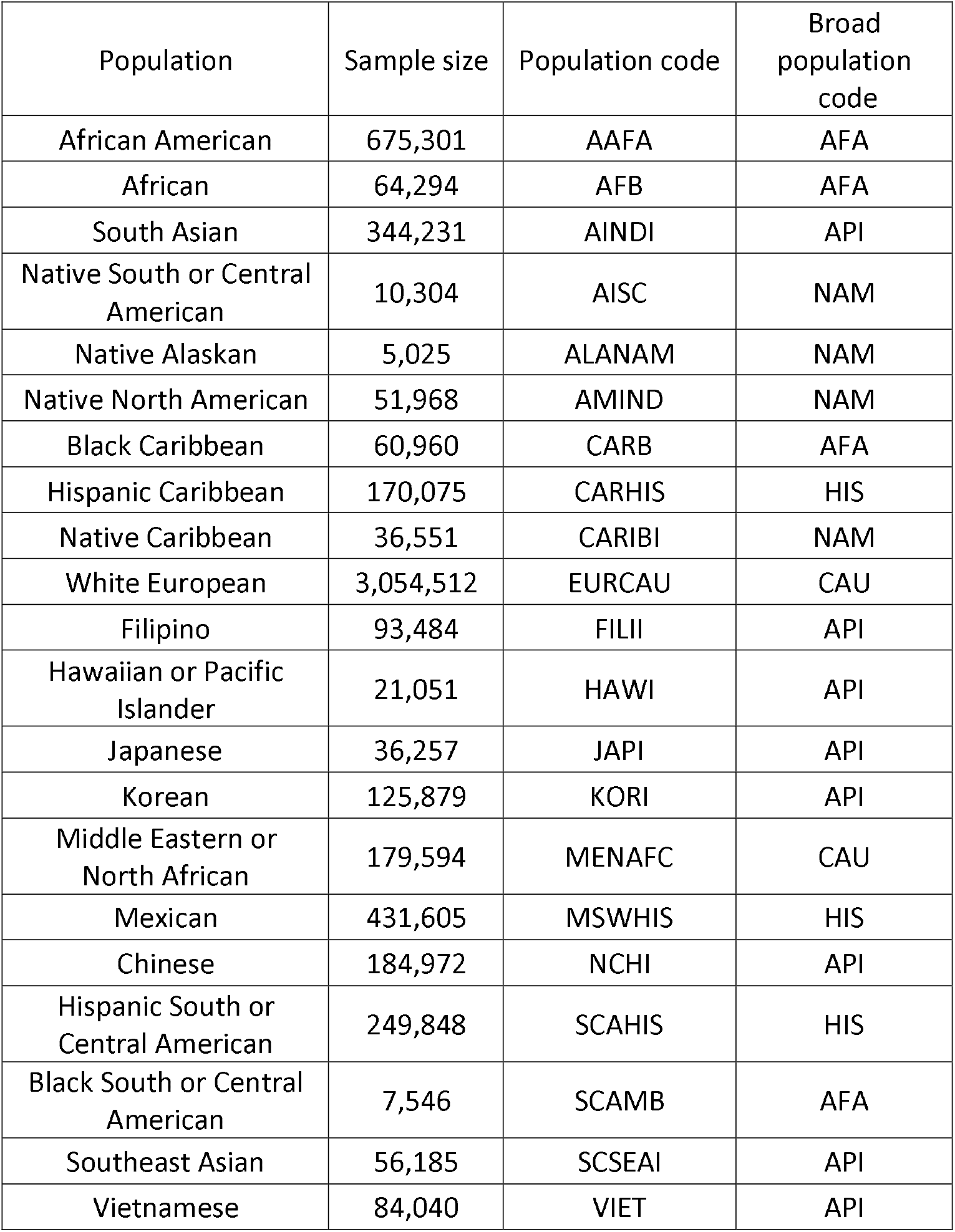
Study sample size by detailed population, including population codes and corresponding broad population categories. This table lists all detailed population groups represented in the study, their sample sizes, and the population codes used in the analyses. Each detailed group is mapped to one of five broad population categories (AFA, API, CAU, HIS, NAM) used for high-level population summaries.

**Table 2.** Broad population sample sizes, including all individuals assigned to each category through detailed population codes or broad-level self-identification. Counts reflect the aggregation of all detailed groups falling under each broad population category, along with individuals for whom only broad population information was available.

| Population | Sample Size | Broad population code |
| --- | --- | --- |
| Black | 910,227 | AFA |
| Asian or Pacific Islander | 1,060,005 | API |
| White | 6,145,749 | CAU |
| Hispanic | 1,439,993 | HIS |
| Native American | 115,108 | NAM |

### HLA Genotyping Methods and Resolution Harmonization

Volunteer members joining the registry in the last decade were typed using NGS technologies with more recent recruits sequenced using a mix of NGS whole gene coverage for HLA class I (5’UTR to 3’UTR) and long range phased exons 2 and 3 for HLA class II or NGS targeted exon (exons 2 and 3 for HLA class I and exon 2 for HLA class II) performed by multiple contracted laboratories that use in-house developed protocols and reagents. There were three prior genotyping strategies included in the cohort: 2010-2015: NGS-based genotyping of *HLA-A, HLA-B, HLA-C, HLA-DRB1, HLA-DRB3, HLA-DRB4, HLA-DRB5*, and *HLA-DQB1* on a short read platform targeting only the ARD exons, 2005-2010: a mixture of SSOP methods with some high-resolution Sanger (SBT) genotyping targeting ARD exons with HLA-C being introduced near the end of this period. Prior to 2005, the genotyping was SSOP using a range of kits dating back to the launch of DNA-based registry genotyping in 1996. The number of probes increased over time, so the resolution is a continuum, but the genes targeted were *HLA-A, HLA-B*, and *HLA-DRB1*.

WHO HLA nomenclature assignments were otherwise standardized to the IPD-IMGT/HLA 3.57.0 database version. To prepare HLA genotyping data for analysis, two-field HLA alleles with identical nucleotide sequence in the antigen recognition domain (ARD - exons 2 and 3 for HLA class I and exon 2 for HLA class II alleles) were combined using a Python library developed for conversion between HLA nomenclature assignments at varying resolutions, py-ard (https://py-ard.org) [33]. The Supplementary Methods section provides more details on historical HLA genotyping strategies, data reporting methods, and the impact of genotyping resolution on haplotype frequency analysis. Ambiguous HLA genotypes can be interpreted in the context of ARD-resolution haplotype frequencies and self-identified population categories of the individual, providing a prediction probability distribution of potential ARD-resolution alleles, haplotypes, and multi-locus genotypes [4, 5, 34]. This probability distribution can be used to measure the level of genotyping ambiguity conditional on population haplotype frequencies using a typing resolution score (TRS) metric. Estimates of the TRS for each studied population were assessed as described in [35] which provide an indication of the level of ambiguity in each studied population over the years. TRS values range from 0 (fully ambiguous genotyping) to 1 (no ambiguity).

### US Population analysis categories

Self-identified race and ethnicity were captured on the NMDP registry member volunteer recruitment form [36] which has changed over time. These self-identified categories are mapped to 21 detailed and five broad populations that are used for matching, frequency estimation, and registry modeling [3, 7, 24]. These population categories have been shown to correlate with HLA better than other systems of self-identification [37, 38] and can be predicted from HLA data at an individual level [6]. These findings support that our US population categories derived from forms data are valuable for key HLA population genetics applications, even though the use of categories has limitations given that human genetic variation is continuous rather than discrete. [24]

### Allele and haplotype frequency estimation from ambiguous genotyping data

We previously implemented an expectation-maximization (EM) algorithm to simultaneously resolve phase and allele-level ambiguity to estimate population haplotype frequencies from targeted HLA genotyping data [23, 24]. To achieve nine-locus haplotype frequency estimates on this dataset of mixed-resolution genotyping, we applied two approaches to limit the number of genotypes without biasing the frequency estimates. First, we identified a minimal set of alleles that could explain all genotypes in a population dataset. Second, we applied a partition-ligation approach using repeated two-gene EM haplotype frequency estimates to build nine-locus haplotype blocks, pruning off very low probability haplotype pairs at each step. Detailed methods on other computational aspects are described in Supplementary Methods (sections 4 and 5).

All HLA genotypes had phase ambiguity because the genomic regions between genes were not sequenced. Our EM algorithm manages varying levels of genotyping resolution, even if the level of genotyping resolution is not independent of the HLA locus (for example there is a higher rate of missing genotyping at genes like *HLA-DPA1* and *HLA-DQA1*) and some alleles were reported with ambiguity represented using multiple allele codes. *HLA-DRB3, HLA-DRB4* and *HLA-DRB5* genes were combined into a single locus following documented and published behavior that they segregate [39-41] and accounting for known associations between them in haplotype estimation [42, 43].

### Profiling of HLA Diversity

#### 1. Asymmetric Linkage Disequilibrium patterns

Asymmetric Linkage Disequilibrium (ALD) among HLA genes was estimated for each population using an ALD method [44, 45] that captures the asymmetry among genes due to a different number of alleles for each HLA locus. ALD provides more information about the LD structure and direction over conventional symmetric LD measures. Unlike symmetric LD statistics, ALD quantifies the conditional dependence of each locus on the other independently, yielding two directed measures per locus pair that reflect the asymmetry in predictive information arising from differences in allelic diversity between loci. For example, knowledge of *HLA-DRB1* polymorphism can inform the prediction of *HLA-DQB1* polymorphism with higher accuracy than the opposite direction since the ALD between both genes is much stronger in the former direction than the latter. The R AsymLD 0.1 package was used for the ALD calculations.

#### 2. Population Clustering and Principal Component Analysis

Multiple population genetics analyses were conducted to compare the US populations at the nine-locus level (*HLA-A, HLA-C, HLA-B, HLA-DRBX, HLA-DRB1, HLA-DQA1, HLA-DQB1, HLA-DPA1, HLA-DPB1*). We use *HLA-DRBX* to refer to the composite locus encompassing the genes *HLA-DRB3, HLA-DRB4*, and *HLA-DRB5* or their absence, treated as mutually exclusive alleles conditional on *HLA-DRB1*.

To visualize population variation, we performed multi-dimensional scaling via Principal Components Analysis (PCA) to summarize frequency difference among populations through dimensionality reduction of the entire nine-locus haplotype frequency distribution of each of the included populations. PCA and corresponding plots were created using the libraries ggplot2 [46], ggfortify [47] and cluster [48] R version 4.2.2 (2022-10-31).

#### 3. Haplotype diversity within populations

To quantify the haplotypic diversity in the studied populations, we sampled 100,000 discrete haplotypes from the entire nine-locus frequency distribution for each broad and detailed population category and depicted their cumulative frequency distributions overlaid on the same plot for all the broad populations: Black, Asian or Pacific Islander, White, Hispanic, and Native American as well as the detailed groups under each of these broad categories. Sampling from the entire distribution accounts for both genetic diversity and sample size effect since inclusion of more samples changes the shape of the haplotype frequency distribution. To adjust for differences in sampling depth among population datasets and to illustrate HLA haplotype diversity on the same scale, we repeated the experiment on equal-sized simulated populations. We simulated five broad and 21 detailed equal sized samples of 50,000 individuals from all the studied groups and repeated the above exercise, therefore excluding the sample size effect and accounting only for genetic diversity of each population. The 50,000-sample size was chosen to mimic the smallest sample size among broad population categories in our dataset (Native Americans).

#### 4. Visualizing Sharing of Common Haplotypes Among Populations

To visualize the degree of overlap in the 100 highest ranked nine-locus haplotypes among the five broad US population categories, an UpSet plot was created. UpSet plots follow a similar concept to Venn diagrams but utilize a bar plot to show the distribution of haplotype counts that are shared among each individual broad population and all observed combinations of populations.

To characterize similarities in haplotype frequency profiles across populations, we performed clustering of the top 100 nine-locus haplotypes using the CLUTO package [49]. Haplotype frequency vectors were row-normalized by population, and pairwise similarity between haplotypes was computed based on their distribution across the 21 populations. A neighbor-joining tree was then constructed to group haplotypes with similar population frequency profiles, enabling identification of clusters representing shared or population-enriched haplotypic patterns.

#### 5. Frequency Comparison with previous studies

To enable comparison with prior HLA haplotype frequency resources, we evaluated concordance between the present dataset and previously published NMDP and international studies across overlapping genes and populations. Comparisons included earlier NMDP datasets at three-locus DRB1∼DQA1∼DQB1 [50], two-locus DPA1∼DPB1 [51], six-locus extended haplotypes combinations [24], as well as full nine-locus classical HLA haplotype datasets from the 17^th^ International HLA and Immunogenetics Workshop[52, 53]. Concordance was quantified using the Renkonen similarity index [54], *If*, which measures the proportion of shared haplotype frequency distributions between datasets while accounting for sampling variance. To further assess external validity of population structure, allele frequency distributions were compared with reference populations from the 1000 Genomes Project [55] using the Renkonen similarity index averaged across nine classical HLA genes, enabling cross-cohort evaluation of population-level genetic similarity. Amino acid level variation for HLA-DP motifs was computed using the R package HLATools [56].

## RESULTS

### Complete US Allele and Haplotype Frequency Datasets are Publicly Available

A comprehensive dataset of estimated allele and haplotype frequencies for all US population categories is publicly available in machine-readable comma-separated value (CSV) format under a Creative Commons license (CC BY-NC-ND 4.0). A trimmed version of the frequency data is also provided in Excel spreadsheet format to support routine use and inspection. The datasets can be downloaded at https://zenodo.org/records/17966993. Population category definitions, sample sizes, and filename population codes are summarized in Table 1 (21 detailed population categories) and Table 2 (5 broad population categories).

### Registry Genotyping Coverage and Resolution Increased Substantially Over Time

Genotyping ambiguity and incomplete locus coverage are inherent challenges in registry-scale HLA data because donor genotypes are accumulated over many years while clinical matching guidelines continue to evolve [28]. To quantify ambiguity in the genotypes used for haplotype frequency estimation, we applied the typing resolution score (TRS) metric [35], which measures ambiguity conditional on population haplotype frequencies. As expected, earlier probe-based genotyping eras showed substantially higher ambiguity. Ambiguity decreased markedly after adoption of sequence-based genotyping platforms beginning in 2012 and decreased further in the most recent recruitment period with implementation of nine-locus typing (Figure 1).

**Figure 1.**
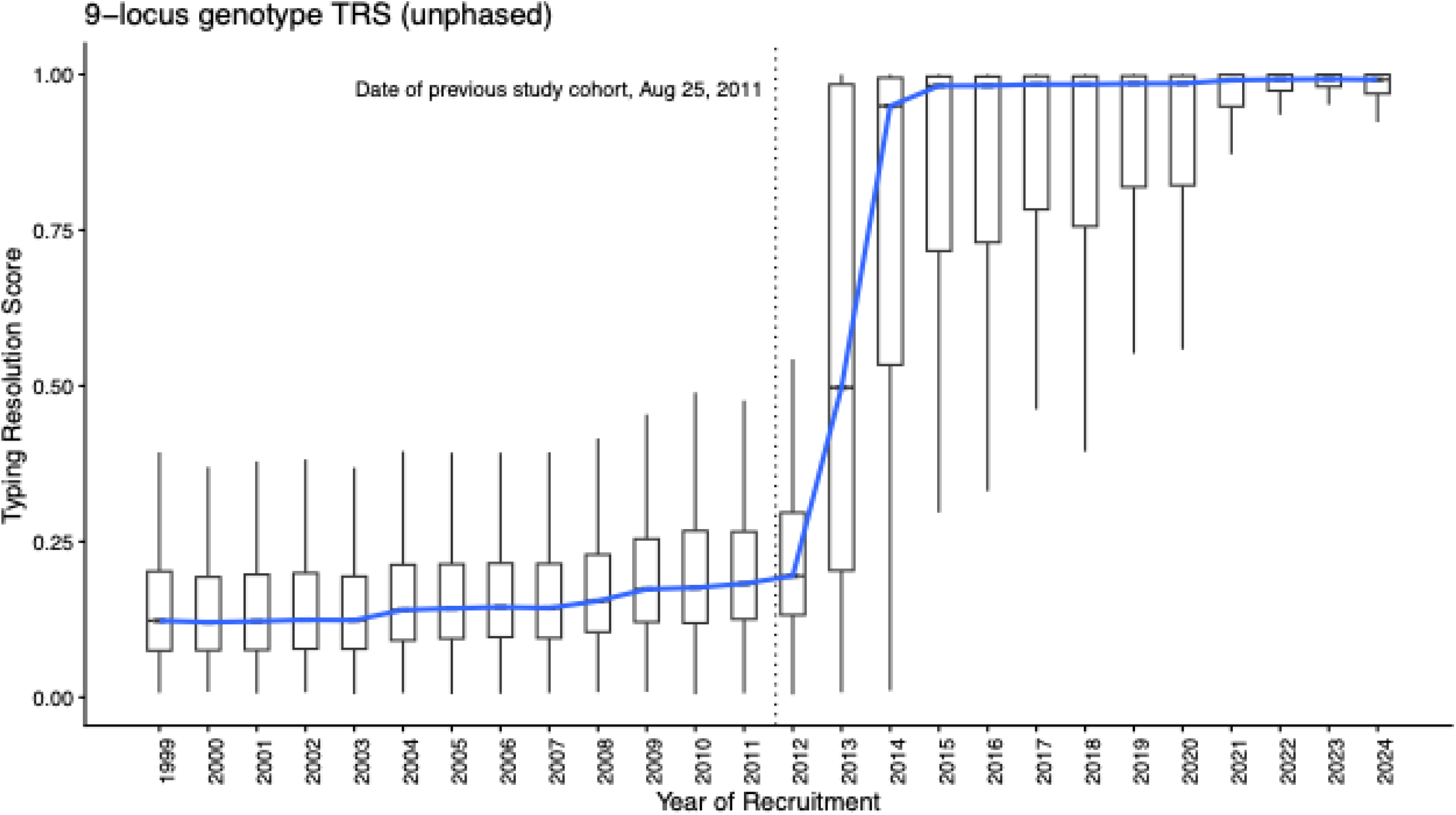
Evolution of Typing Resolution Score (TRS) among NMDP donors. Nine-locus Typing Resolution Score (TRS) from DNA-typed NMDP donors over time. The figure illustrates the progressive improvement in molecular HLA typing resolution across recruitment years, reflecting changes in laboratory technologies and typing policies.

Genotyping coverage and resolution also increased over time across loci. All donors included in this study were DNA-typed at minimum for *HLA-A, HLA-B*, and *HLA-DRB1*. The fraction of donors typed at *HLA-C* increased from 25% to 64%, and typing at *HLA-DQB1* increased from 6% to 54%. Among the additional HLA class II genes included in this analysis, 11% of donors were typed at *HLA-DQA1* and *HLA-DPA1*, and 42% at *HLA-DPB1* (Table 3). Nearly all *HLA-DQA1, HLA-DQB1, HLA-DPA1*, and *HLA-DPB1* genotypes were reported at high resolution, whereas a smaller fraction of *HLA-A, HLA-B, HLA-C*, and *HLA-DRB1* genotypes were high resolution because these loci were routinely typed in earlier eras under clinical matching guidelines before next-generation sequencing became standard practice.

**Table 3.**
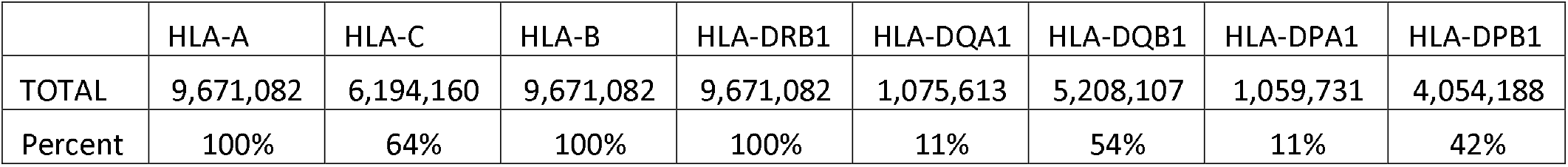
Number of individuals genotyped for each HLA locus included in the study. The overall sample size across all included NMDP broad population categories is provided for each respective locus in the TOTAL row, with the fraction of individuals with genotyping at the locus provided in the “Percent” row. Because eligibility for inclusion required typing at HLA-A, HLA-B, and HLA-DRB1, these loci have identical sample sizes and represent 100% of the cohort. Differences in typing rates for other loci reflect historical changes in laboratory testing policies and platform adoption over time. HLA-DRB3, -DRB4, and -DRB5 are not listed because laboratories typically do not report the absence of these genes; therefore, it is not possible to distinguish between “not typed” and “gene not present” across all individuals.

### Allele Frequencies Were Largely Consistent with the Common, Intermediate, and Well Documented (CIWD) Allele Catalog

We profiled the distribution of HLA allele frequencies from this dataset at the high-resolution level after antigen recognition domain (ARD) within the Common, Intermediate, Well Documented, and Rare categories within 6 CIWD broad population categories in Figure 2. Note the CIWD study [57] separated the White broad population category used in this study into two categories: European (EURO) and Middle Eastern and North African (MENA). For each population, the median frequency of the allele frequency distributions showed stepwise decreases from Common to Rare. With standardization of typing resolution in this frequency study and larger sample size relative to CIWD, we were able to identify that some alleles have a population frequency that could have been a better fit in other CIWD categories. Because the threshold for assigning alleles to the CIWD Well Documented category was based on counts of reported HLA genotypes, CIWD Rare alleles had the lowest allele frequency distribution within the European category because most donors in the CIWD cohort were European.

**Figure 2.**
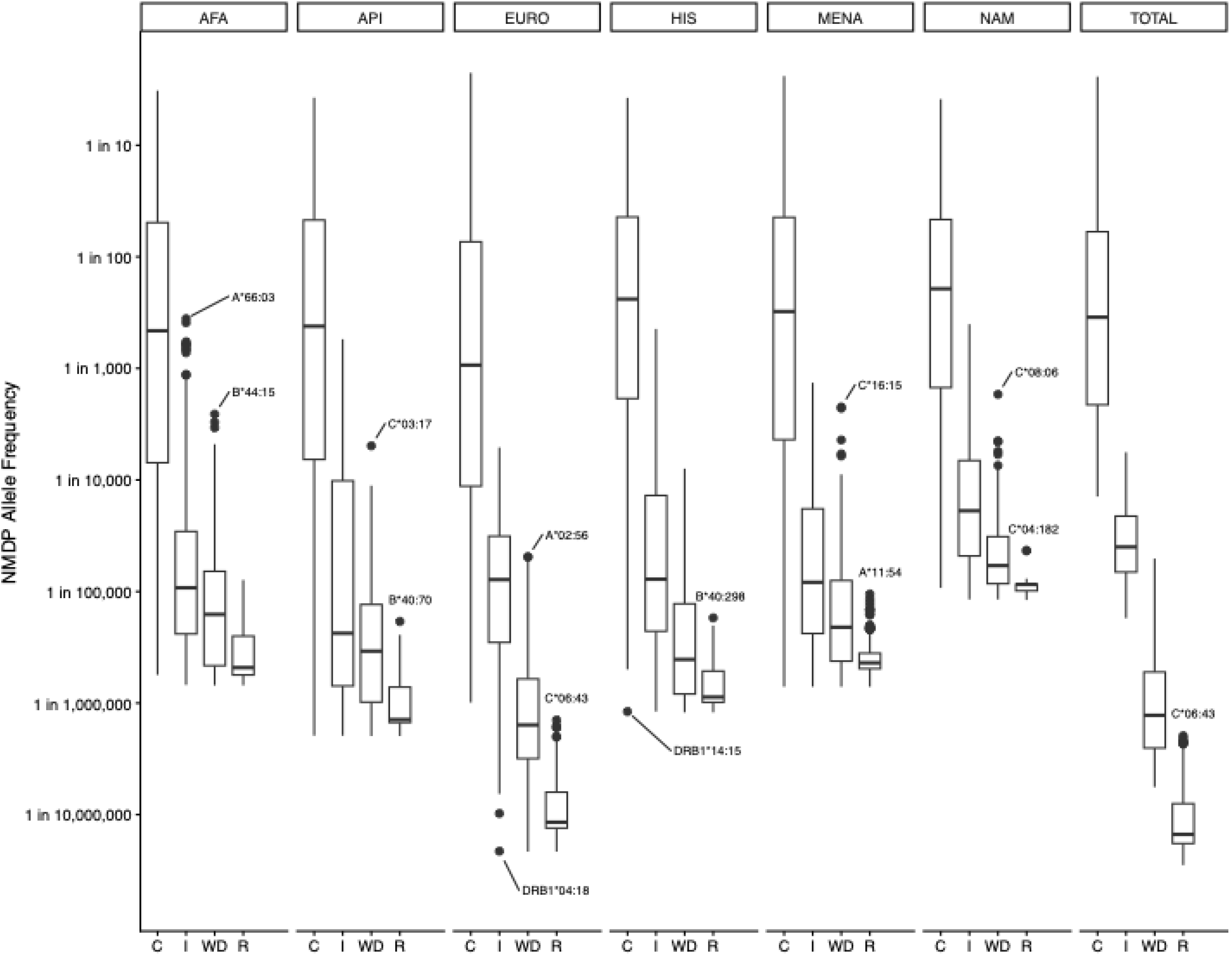
Comparison of HLA allele frequencies from the NMDP registry for the loci HLA-A, -B, -C, -DRB1, -DQB1, and -DPB1 with the CIWD catalog. CIWD data were reduced to ARD equivalence to enable direct comparison. The categories C, I, WD, and R correspond to the CIWD classifications *Common, Intermediate, Well Documented*, and *Rare*. Population groups were aligned with those used in the CIWD study: AFA (African/African American), API (Asian/Pacific Islander), EURO (European/European descent), MENA (Middle East/North Coast of Africa), HIS (South or Central America/Hispanic/Latino), and NAM (Native American). Boxplots show the distribution of allele frequencies within each CIWD category. Alleles falling outside ±1.5× the interquartile range (IQR) are considered outliers, and the most extreme outlier allele is labeled.

### Common Classical HLA Haplotypes Show High Population Specificity

Population distribution of the top 100 most frequent nine-locus HLA haplotypes showed pronounced genetic structure across US population categories (Supplementary Material 1; Figure 5). The majority of high-frequency haplotypes were population-specific or restricted to a small number of related populations, indicating strong differentiation in haplotype frequency among groups. Most top-ranked haplotypes fell predominantly within Asian or Pacific Islander or Black populations, while a separate set was primarily concentrated within White, Hispanic, and Native American populations, which exhibited greater mutual sharing consistent with admixture and shared ancestry. Within the Asian or Pacific Islander category, most top haplotypes were confined to one or two detailed subpopulations, demonstrating substantial within-category heterogeneity. In contrast, widespread sharing across all five broad populations was rare: only three haplotypes appeared in the top-100 lists for every population (Supplementary Figure 1). Together, these frequency sharing patterns indicate that common high-resolution HLA haplotypes are typically population-enriched, with limited cross-population overlap at the highest frequency ranks.

### HLA-DPA1, HLA-DPB1, and HLA-DQA1 Display Limited Allelic Diversity Compared to Other Classical HLA Loci but Strong Population Stratification

Allele frequency distributions differed substantially across populations and loci, with the strongest population differentiation observed at *HLA-DPA1, HLA-DPB1*, and *HLA-DQA1*, genes that have not previously been characterized at this scale across US populations (Supplementary Material 1). These loci show relatively low allelic complexity but pronounced shifts in dominant alleles between broad population categories. For example, *HLA-DPA1* is largely dominated by a small number of alleles whose rank order changes by population group, with *HLA-DPA1*01:03* most frequent in White, Native American, and Hispanic populations, while *HLA-DPA1*02:02* is enriched in many Asian or Pacific Islander populations and *HLA-DPA1*03:01* occurs at appreciable frequency primarily in Black populations. A similar population-stratified pattern is observed at *HLA-DPB1*, where *HLA-DPB1*04:01* is most common in White, Native American, and Hispanic populations, *HLA-DPB1*05:01* is enriched in Asian or Pacific Islander populations, and *HLA-DPB1*01:01* shows higher prevalence in Black populations.

*HLA-DQA1* likewise exhibits a constrained allele diversity dominated by three alleles across populations, but with marked differences in relative frequencies by group, reinforcing population structure at HLA class II loci beyond *HLA-DRB1* and *HLA-DQB1*. In contrast, classical HLA class I loci (*HLA-A, -B, -C)* and *HLA-DRB1* show higher allelic diversity with broader sharing of common alleles across populations, although several alleles remain population-enriched (Supplementary Material 1). Frequencies of *HLA-DRB3, -DRB4*, and -*DRB5* alleles reflect known linkage with *HLA-DRB1*, with *HLA-DRB4*01:01* predominating due to its association with common *HLA-DRB1*04* and *HLA-DRB1*07* bearing haplotypes.

### Nine-Locus Classical HLA Haplotypes Reveal Distinct Population Genetic Structure

We used principal component analysis to summarize differences in nine-locus HLA haplotype frequency distributions among the studied populations. The ordering of principal components reflects which haplotypes contribute the greatest variance across populations. The first two components captured large frequency differences among Asian or Pacific Islander populations, resulting in broader dispersion of these groups along PC1 and PC2 (Figure 3). In contrast, detailed population groups within the Black, Hispanic, White, and Native American broad categories clustered more tightly in this space, indicating greater similarity in their haplotype frequency distributions.

**Figure 3.**
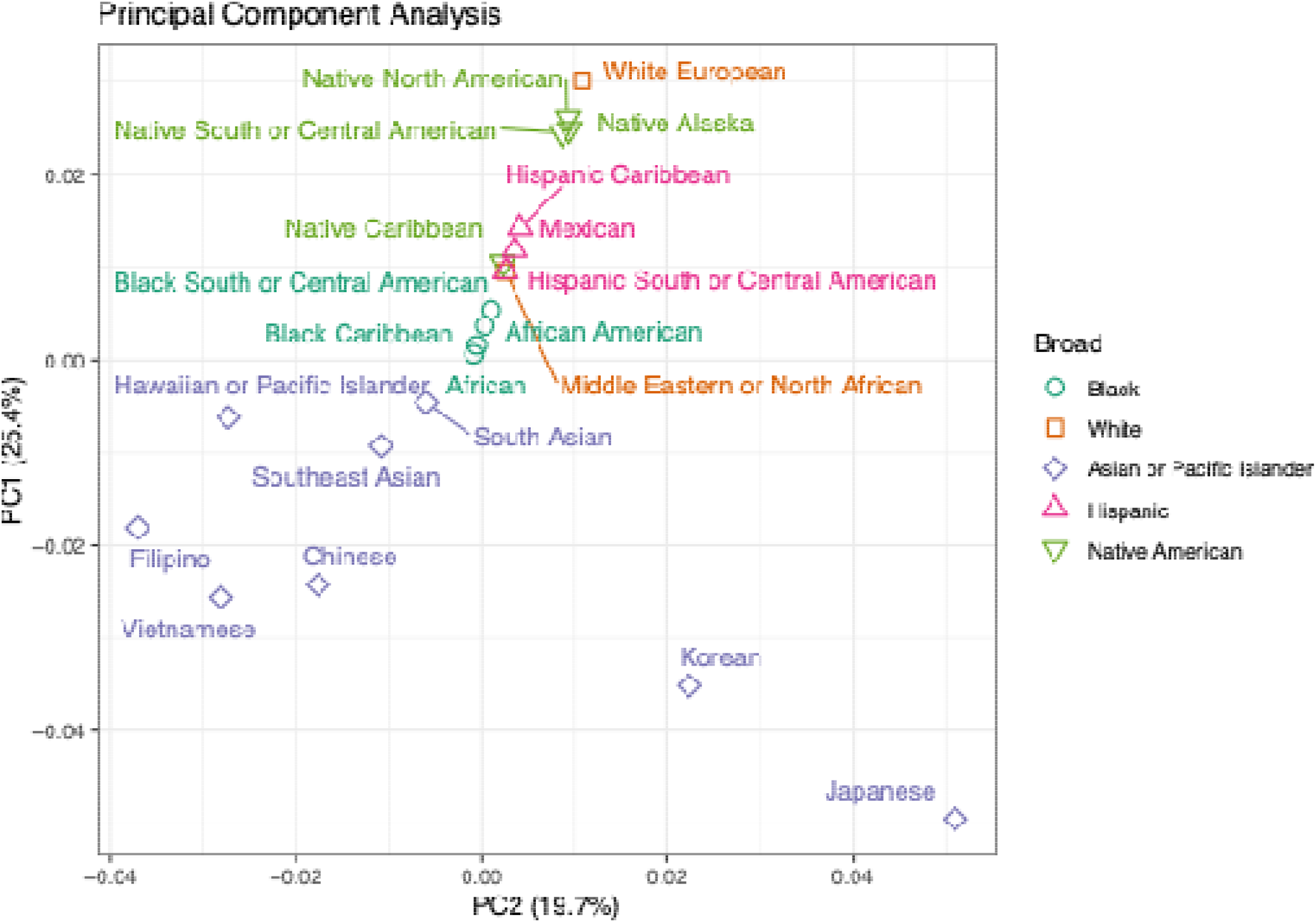
Principal Component Analysis (PCA) of 9-locus haplotype frequencies. PCA performed on population-level 9-locus HLA haplotype frequencies. The first two principal components explain more than 45% of the total variance. Points are colored by broad population group, demonstrating the major axes of genetic structure represented in the donor registry.

Haplotypes that are common across multiple populations—particularly those shared broadly among European and African ancestry groups—contribute less to the variance captured by the first principal components and therefore have lower factor loadings in PC1 and PC2. Population structure associated with these more widely shared haplotypes is instead reflected in later principal components that capture smaller, more broadly distributed frequency differences across the remaining population categories.

### Black Populations Exhibit the Greatest Haplotypic Diversity Across All Analyses

We evaluated nine-locus haplotype diversity across the studied US populations by examining cumulative frequency distributions under two complementary sampling frameworks (Figure 4). In the first analysis, haplotypes were sampled directly from the estimated population-specific frequency distributions, thereby reflecting both underlying genetic diversity and differences in registry sample size. In the second analysis, equal-sized simulated populations were generated to isolate differences in haplotypic diversity independent of sampling depth.

**Figure 4.**
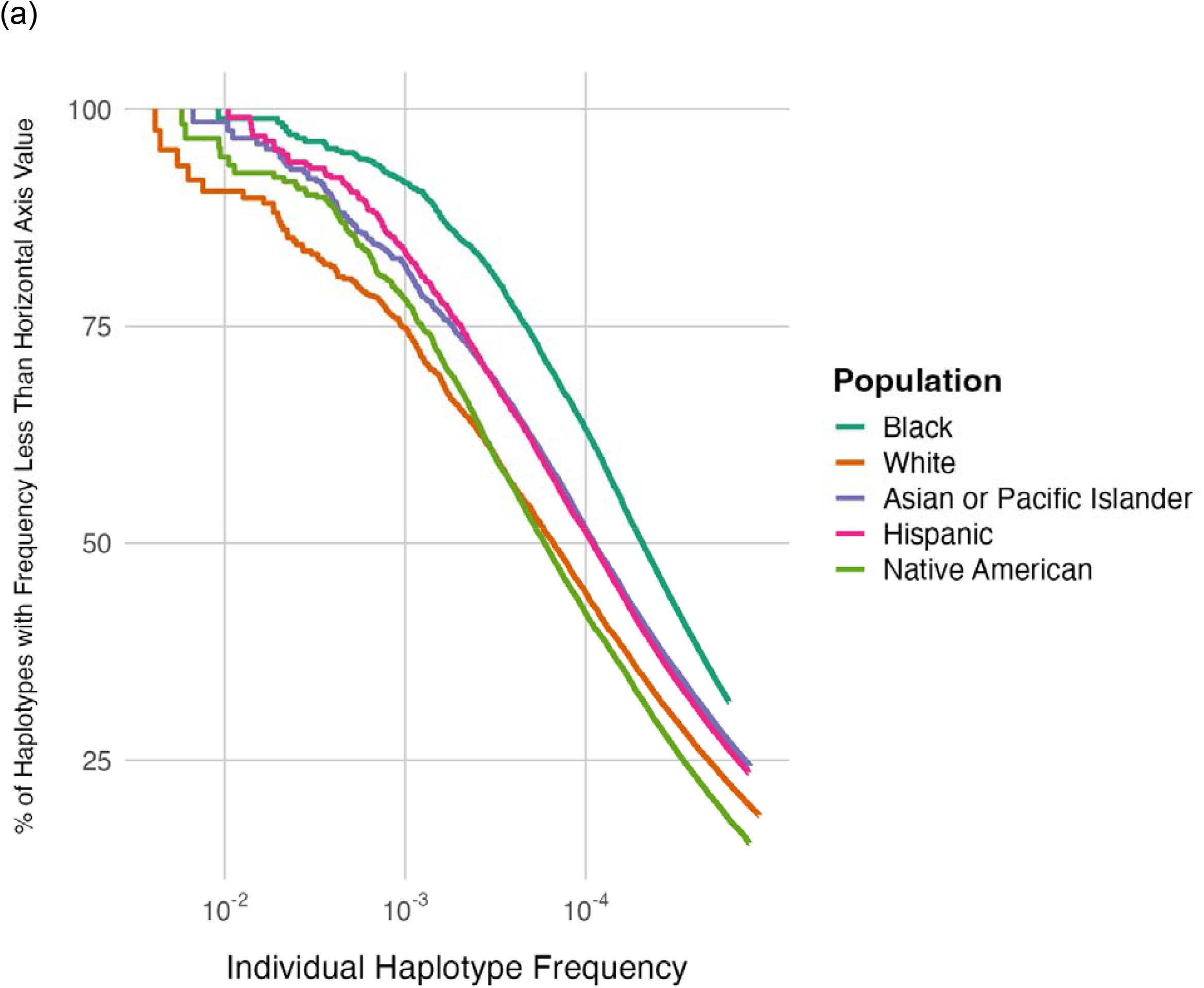
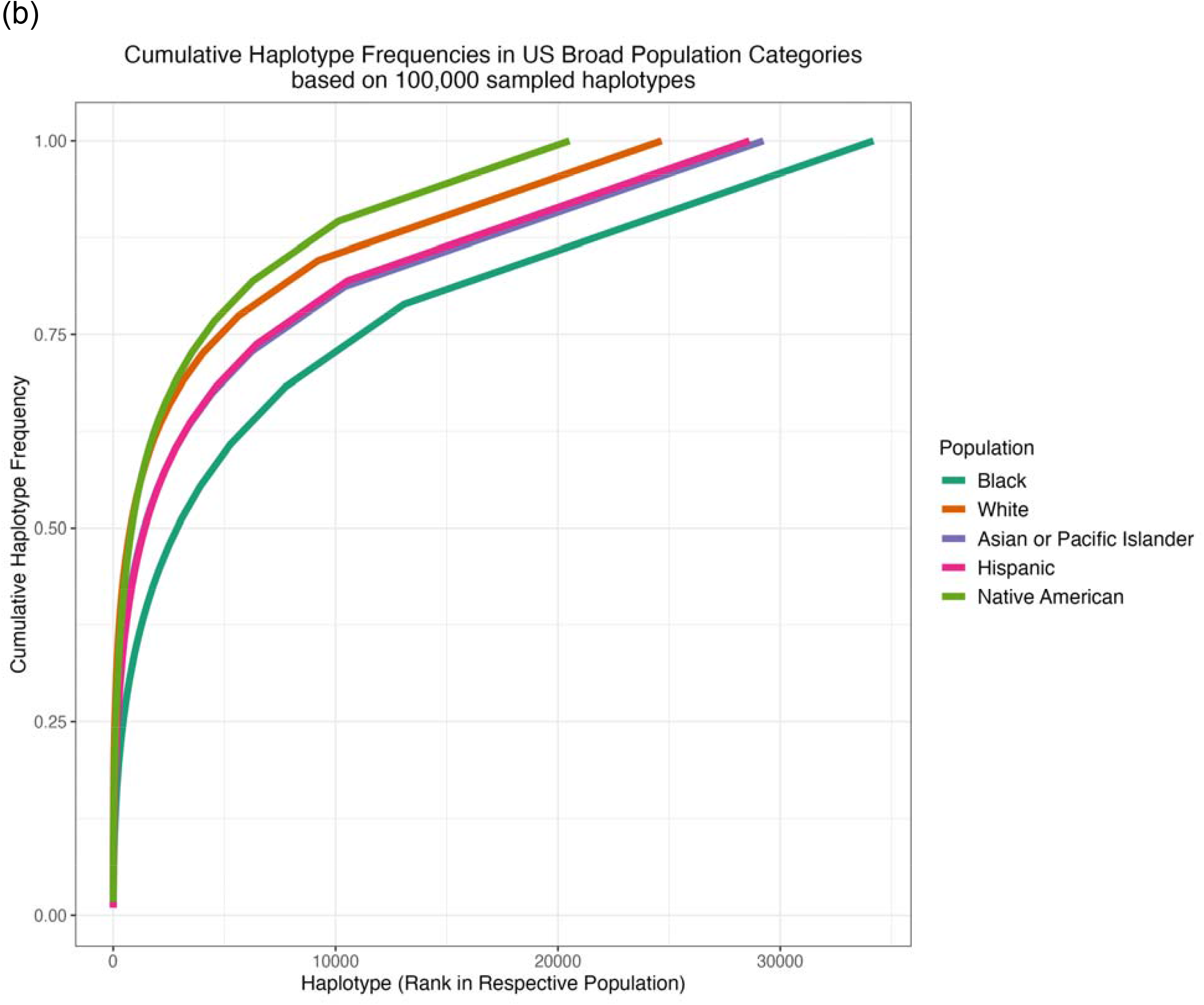
Cumulative HLA haplotype frequency distributions among populations. **(a)** Unequal sample size using the complete haplotype frequency estimates - Cumulative haplotype frequency plot showing the portion of the population, on the Y-axis, with frequency less than the frequency value on the X-axis for the top 10,000 haplotypes. Curves further to the left are less diverse with fewer high frequency haplotypes accounting for a larger fraction of the population. **(b)** Equal sample size - Comparative coverage across detailed and broad population groups, illustrating variability in match likelihood across populations given an equal sample of 100,000 sampled haplotypes (50,000 genotypes after realization from high resolution probability distribution) per population. The knees in the righthand part of the curves indicate singleton sampled haplotypes.

When sampling from the full population-specific frequency distributions, Black populations exhibited the greatest haplotypic diversity. Relative to other broad population categories, a substantially larger number of distinct haplotypes were required to account for a given proportion of cumulative frequency. This pattern reflects a flatter haplotype frequency distribution characterized by a long tail of low-frequency haplotypes. In contrast, Native American populations showed the most concentrated haplotype frequency distribution, followed by White populations. Hispanic and Asian or Pacific Islander populations displayed intermediate diversity patterns.

To distinguish genetic diversity from the effects of unequal sample sizes across registry populations, we repeated the analysis using simulated populations of equal size. Under this normalization, Black populations remained the most diverse, confirming that the broader haplotype distribution is not solely a consequence of larger representation in the registry. Asian or Pacific Islander and Hispanic populations also showed relatively high haplotypic diversity, whereas White and Native American populations exhibited more concentrated haplotype frequency distributions. Notably, the apparent diversity observed in White populations under the full-sample analysis decreased after equal-size normalization, indicating that part of the observed haplotype breadth in that group reflects the large cohort size rather than underlying genetic diversity alone.

### HLA-DRB1 Is the Strongest Directional Predictor Across the HLA class II Region

To characterize directional relationships among HLA loci, we estimated asymmetric linkage disequilibrium (ALD) across the nine classical HLA genes for each studied population (Figure 5; Supplementary Materials 2). Unlike conventional symmetric linkage disequilibrium measures, ALD quantifies whether knowledge of one locus is more informative for predicting another than vice versa, which is particularly useful for loci that differ in allelic diversity and typing completeness.

**Figure 5.**
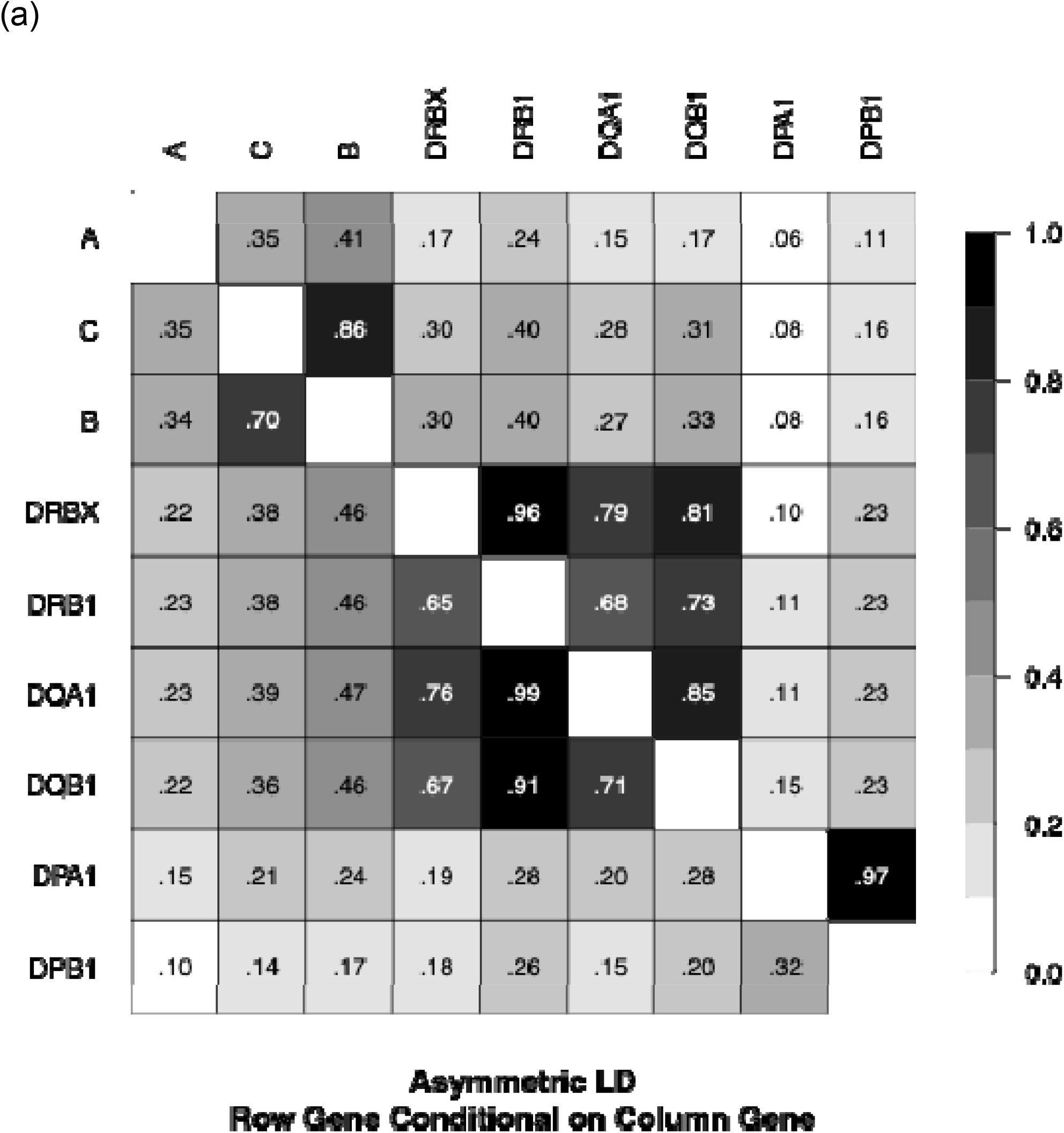
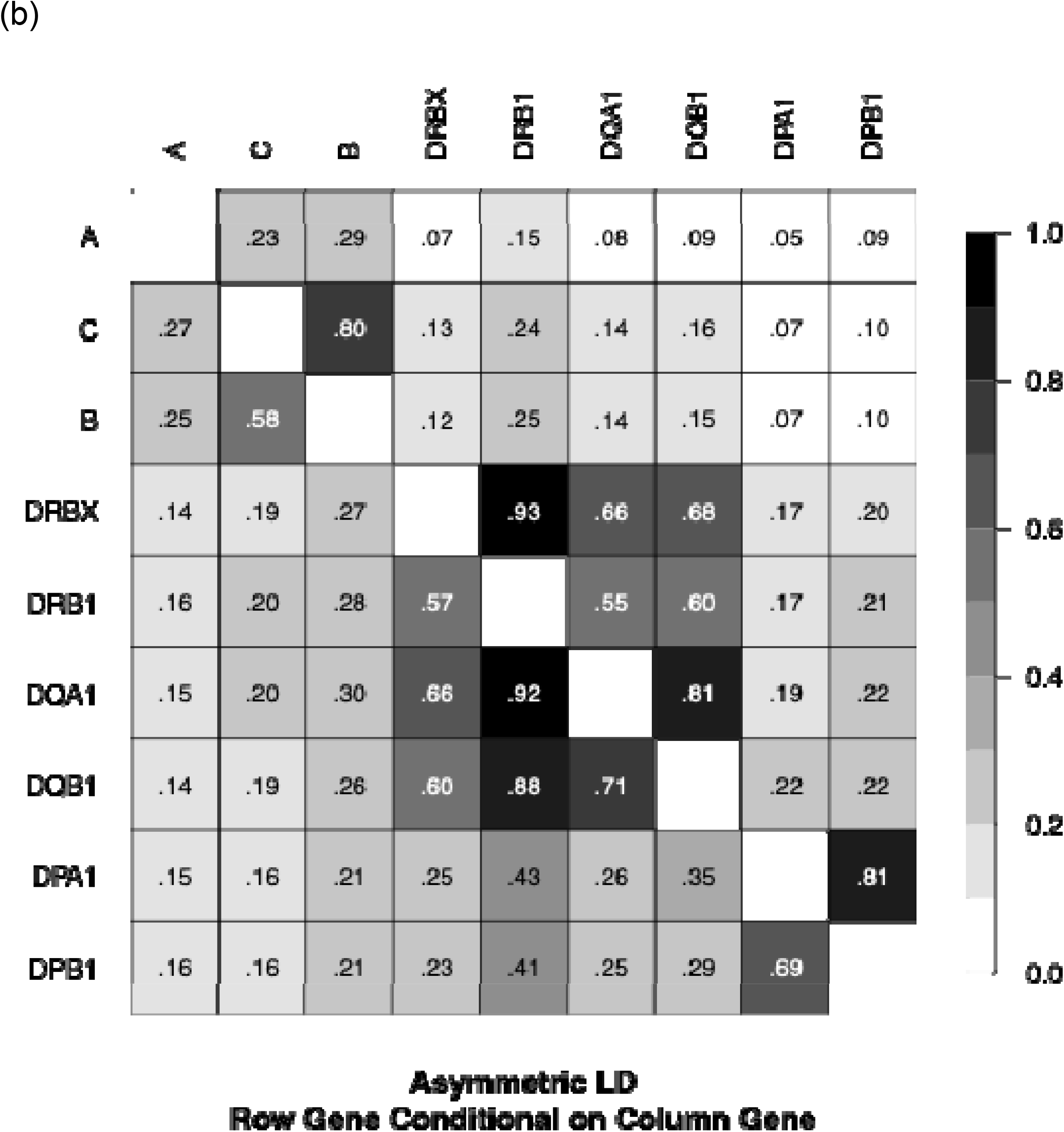
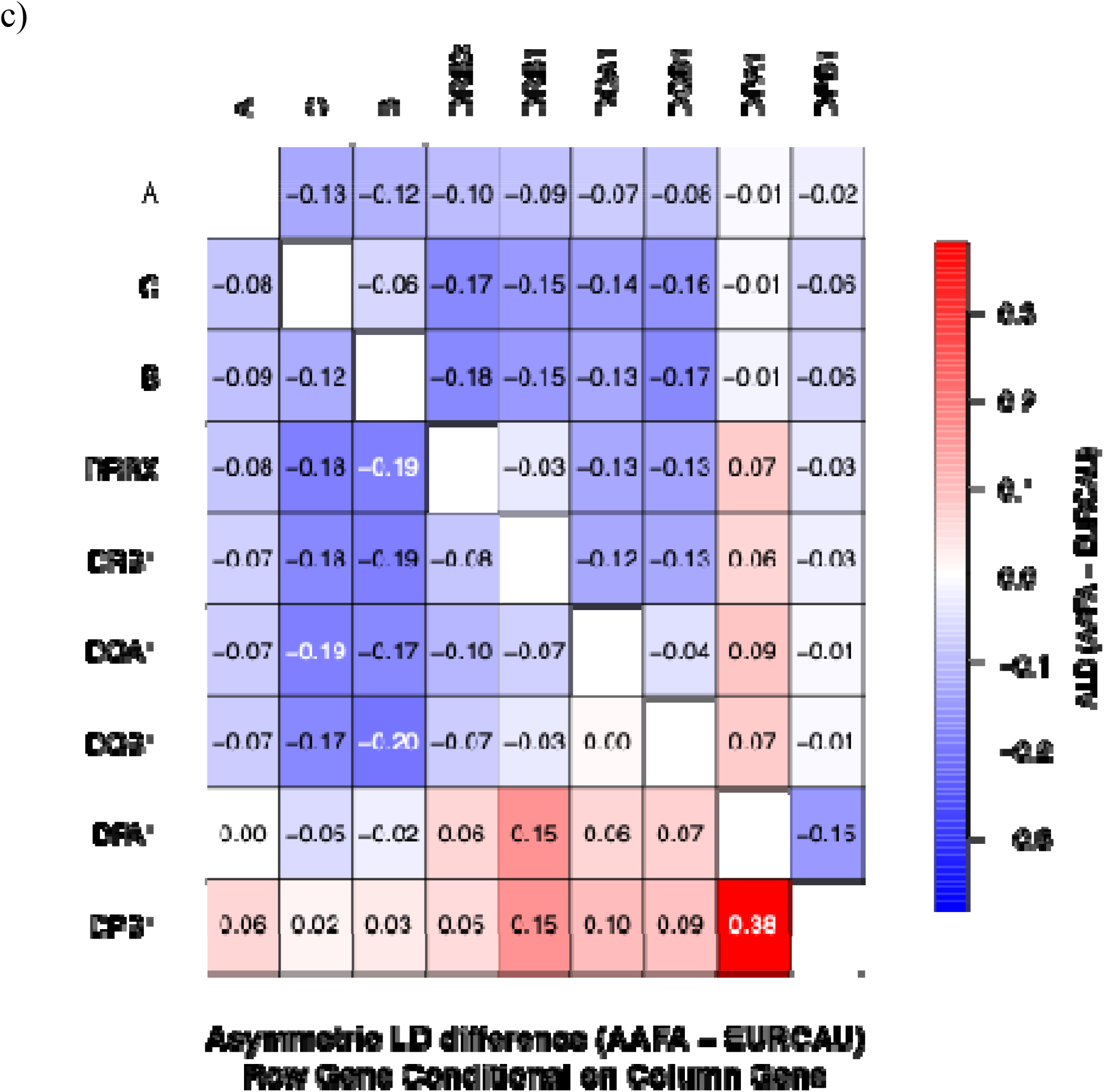
Asymmetric Linkage Disequilibrium (ALD) for major populations. ALD estimates based on 9-locus HLA haplotypes. The direction of the association is read as the row gene conditional on the column. The two panels: (a) White European (EURCAU). (b) African American (AAFA), highlight contrasts in LD these populations. (c) Difference in LD values (African American – White European). Note: substantially higher LD is observed in both populations from DRB1 (column) to DQA1 (row) compared with DQB1 (column) to DQA1 (row) even though DQA1 and DQB1 form a heterodimer. The data for all populations is provided in Supplemental Materials 2.

Across populations, the strongest directional associations were observed within the HLA class II region. In particular, *HLA-DRB1* showed strong predictive relationships with the neighboring loci *HLA-DQA1, HLA-DQB1*, and the composite *HLA-DRB3/-DRB4/-DRB5* locus. The strongest directional dependence was consistently observed from *HLA-DRB1* to *HLA-DQA1*, followed by *HLA-DRB1* to *HLA-DRB3/-DRB4/-DRB5* and *HLA-DRB1* to *HLA-DQB1*. These results indicate that knowledge of DRB1 substantially constrains the expected allelic states at adjacent HLA class II loci.

Interestingly, the directional association from *HLA-DRB1* to *HLA-DQA1* exceeded that from *HLA-DQB1* to *HLA-DQA1*, despite the fact that the *HLA-DQA1* and *HLA-DQB1* gene products form the heterodimeric HLA-DQ molecule. This observation highlights that population-level haplotypic structure across the MHC reflects the evolutionary history of extended haplotypes rather than simply protein pairing relationships. We analyzed the frequencies of HLA-DQ Group 1 haplotypes (containing HLA-DQA1*02/03/04/05/06) and HLA-DQ Group 2 (containing HLA-DQA1*01) haplotypes among populations [58], finding balanced frequencies in White populations, but up to 70.9% frequency of HLA-DQ Group 1 in Hispanic populations (Supplementary Table 1).

While the overall ALD structure was broadly similar across populations, directional associations were generally weaker in African Americans than in White Europeans (Figure 5c). Across most loci, lower ALD values were observed in African American populations, consistent with the greater haplotypic diversity and reduced long-range linkage disequilibrium typically observed in populations with deeper ancestral diversity. A notable exception involved the HLA-DP region, where stronger directional associations between *HLA-DPB1* and other HLA loci were observed in the African American population relative to the White European population, suggesting that LD patterns involving *HLA-DPB1* may vary substantially across populations. We analyzed a previously described HLA-DP amino acid level motif comprising *HLA-DPA1* position 31 and *HLA-DPB1* positions 85-87, which was found to be in complete linkage disequilibrium in the White population in a previous NMDP study [51], finding that Asian populations had far lower linkage disequilibrium (Supplementary Table 2).

These results demonstrate that the nine-locus haplotype framework captures strong directional structure across the HLA region, particularly within the HLA class II loci, and helps explain the effectiveness of haplotype-based imputation methods for predicting missing or ambiguous HLA genotypes.

### Population-Specific Haplotypes Predominate, with Admixture Reflected in White– Hispanic–Native American Sharing

The UpSet plot of the 100 highest ranked 9-locus haplotypes among the five broad US population categories (Figure 6) showed strong population specificity of common haplotypes in Asian or Pacific Islander and Black populations, where 90 of the top-100 Asian or Pacific Islander haplotypes were not observed among the top 100 haplotypes of any other broad US population, and likewise 86 of the top 100 for Black. For White, Hispanic, and Native American populations, 39 of the top 100 haplotypes in each population were private. These three populations had the highest degree of haplotype sharing between them, as 55 top-100 haplotypes were shared among two or three of these populations, which is consistent with the large amount of population admixture between these three populations. Nine top-100 haplotypes were shared among four populations, Black, White, Hispanic, and Native American, with most of these haplotypes being very common in Whites, also indicating admixture from Europeans into these other populations. Sharing of top-100 haplotypes between other broad population combinations was far less frequent. Detailed information on the frequency and rank of top-100 haplotypes is available in Supplementary Materials 3.

**Figure 6.**
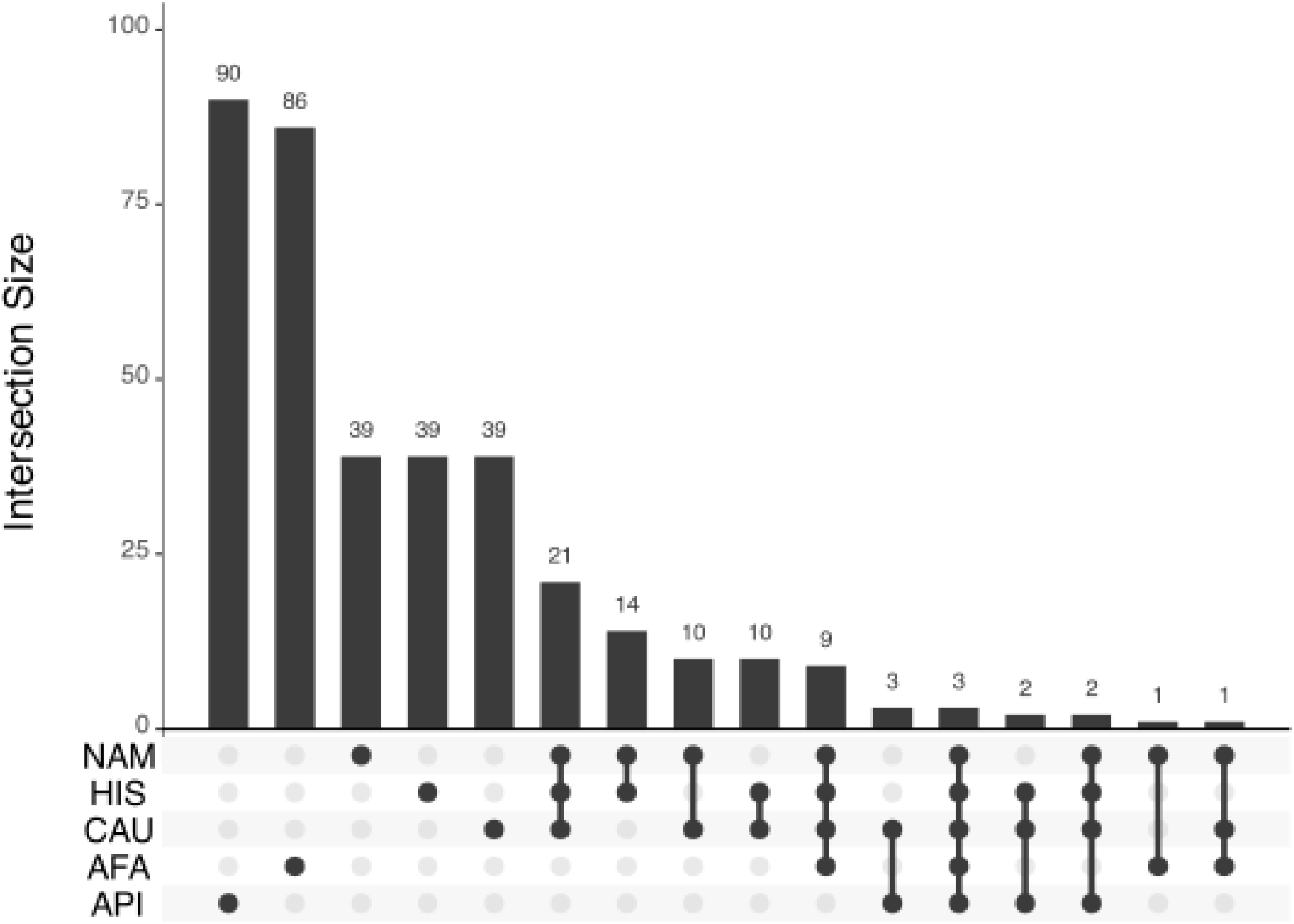
Shared 9-locus haplotype patterns across populations (UpSet plot). UpSet plot showing intersections of HLA alleles or haplotypes shared across populations. The plot quantifies the extent of overlap and identifies population-specific and widely shared genetic features within the registry. The majority of the 100 more frequent 9-locus HLA haplotypes are private to one population group with only 3 haplotypes observed in the top 100 in all 5 population groups.

### Nine-Locus Haplotype Clustering Reveals Population Structure

Clustering of the top 100 nine-locus haplotypes revealed distinct groupings corresponding to major population categories, with haplotypes predominantly clustering according to their highest-frequency populations. Haplotypes common in Asian or Pacific Islander and Black populations formed largely separate clusters, while haplotypes enriched in White, Hispanic, and Native American populations showed greater intermixing, reflecting shared ancestry and admixture. These clustering patterns are consistent with results from principal component analysis and haplotype sharing analyses, reinforcing the presence of strong population structure alongside varying degrees of cross-population overlap.

### Frequency distributions had high similarity with all previous studies

Haplotype frequency estimates from the current dataset showed high concordance with previously published NMDP studies across all major populations. Six-locus comparisons with prior NMDP data yielded *If* values typically ranging from approximately 82% to 90% for African American, Asian or Pacific Islander, European American, and Hispanic populations, with somewhat lower concordance observed in Native American populations, reflecting smaller sample sizes and greater heterogeneity (Supplementary Table 3). Concordance with earlier HLA class II–focused NMDP studies was similarly high (approximately 90% or greater), indicating consistency of haplotype structure despite substantial increases in sample size and locus coverage (Supplementary Table 4). Comparisons with International HLA and Immunogenetics Workshop datasets demonstrated lower overlap, particularly for nine-locus haplotypes, largely due to truncation of frequency distributions in those datasets, which capture only a subset of common haplotypes (Supplementary Table 5). Population similarity analysis with the 1000 Genomes Project showed strong correspondence in overall population structure, with highest similarity observed between genetically matched population groups and expected clustering patterns across continental ancestries (Supplementary Figure 2). Visualizing comparing the 40 most common haplotypes in this dataset with the previous NMDP US frequency estimates published in 2013, refinement of haplotype frequency estimates was observed especially for phasing of the *HLA-DQB1* locus on extended haplotypes. We provide an illustrative example where the greatest frequency difference between Asian or Pacific Islander datasets was observed for a pair of recombinant six-locus extended haplotypes with differences only at the *HLA-DQB1* locus (Supplementary Figure 3). While only 6% of donors were typed for *HLA-DQB1* in the US 2013 dataset, genotyping coverage of *HLA-DQB1* increased to *54%* in this dataset, much of it at high resolution. Inclusion of *HLA-DQA1* likely also greatly improved ability to correctly phase extended haplotypes because of its high LD with other HLA class II loci.

## DISCUSSION

This study provides the most comprehensive high-resolution antigen recognition domain level HLA haplotype frequency resource for US populations to date, derived from nearly ten million volunteer donors in the NMDP registry and encompassing all nine classical HLA loci. The analysis advances prior NMDP frequency resources in three important ways. First, it extends haplotype frequency estimation from earlier three- and six-locus frameworks to nine loci, incorporating the class II genes *HLA-DQA1, HLA-DPA1*, and *HLA-DPB1* that have increasingly been included in donor typing and transplant outcome studies. Second, the analysis leverages a greatly expanded dataset with substantially improved locus coverage and typing resolution compared with previous registry-based frequency estimates. Third, the computational framework used here enables robust estimation of multi-locus haplotype frequencies from mixed-resolution and partially ambiguous HLA genotyping data at a scale that was previously computationally prohibitive.

Several population-genetic features of the HLA system are clearly evident in this expanded dataset. As previously observed in smaller datasets, HLA haplotype frequency distributions follow a heavy-tailed structure in which a small number of haplotypes occur at relatively high frequency while a long tail of rare haplotypes contributes substantially to overall diversity. Despite this long tail, the most frequent haplotypes show strong population specificity. Across the five broad US population groups and the 21 detailed populations examined here, most high-frequency haplotypes are restricted to a limited set of populations, reflecting both historical divergence of ancestral populations and subsequent admixture patterns within the United States.

Population comparisons continue to reveal clear genetic structure across the HLA region. Detailed population groups cluster consistently within their respective broad population categories across multiple analyses, including principal component analysis, haplotype clustering, and linkage disequilibrium patterns. In particular, Black and Asian or Pacific Islander populations form distinct clusters, while Hispanic, White, and Native American populations show greater overlap, consistent with known patterns of admixture. These patterns reflect the persistence of ancestral haplotypes across generations even in the context of increasing demographic mixing. Notably, among the thousands of nine-locus haplotypes estimated across the 21 detailed populations, only a small number are shared across all groups, highlighting the extensive haplotypic diversity present within the US population. The high concordance with prior NMDP datasets indicates that expansion to a nine-locus framework preserves established haplotypic structure while having more complete typing data improved. resolution of rare and population-specific variation. Lower overlap with International HLA and Immunogenetics Workshop datasets reflects differences in haplotype representation, and strong agreement with 1000 Genomes Project populations supports the practical usage of registry-based population categories from race/ethnicity questionnaires while highlighting effects of admixture and sampling.

The expanded nine-locus haplotype frequency framework has several practical implications. In transplantation, these frequencies improve the accuracy of predicting high resolution ARD-level HLA genotypes and haplotypes from ambiguous or partially typed donor and patient data. Integration of these frequencies into matching algorithms such as HapLogic^SM^ enables improved prediction of donor–recipient compatibility, particularly for loci that until recently have not been universally typed in registry donors, including *HLA-DQA1* and *HLA-DPA1*. The resulting improvements in genotype inference and match likelihood estimation can enhance the efficiency of donor search processes and improve matching strategies for patients lacking fully typed donors.

Beyond transplantation, population-specific haplotype frequency data support several areas of biomedical research. The expanded coverage of class II loci facilitates improved analyses of linkage disequilibrium and haplotype structure in studies of immune-mediated disease. In addition, the characterization of allele and haplotype distributions across US populations informs the design of immunotherapies and vaccines by enabling population-level estimates of HLA-mediated epitope presentation. Finally, these data provide a valuable reference for modeling donor registry composition and evaluating strategies to improve equitable access to transplantation across diverse populations.

The Common, Intermediate, and Well Documented (CIWD) catalog [57] is an important community reference used to guide assay design and policies for resolving HLA genotyping ambiguities. However, direct comparison between CIWD classifications and the allele frequency estimates reported here is limited by methodological differences. The frequencies in this study were estimated using a unified maximum-likelihood framework in which all genotypes were harmonized to a consistent antigen recognition domain (ARD) resolution and interpreted relative to a single reference database version. In contrast, the CIWD catalog aggregates genotyping observations reported to multiple registries across heterogeneous sequencing platforms, typing resolutions, and analysis pipelines. Because some CIWD source datasets retain unresolved ARD-level ambiguities, the resulting allele counts are not directly comparable to the ARD-standardized frequency estimates presented here. Future comparisons will benefit from fully phased full-gene sequencing datasets analyzed under consistent laboratory and informatics pipelines, which will provide a stronger foundation for revisiting CIWD classifications at higher levels of resolution[59-61].

This study has several limitations. First, because the underlying data derive from targeted HLA genotyping assays rather than whole-MHC sequencing, haplotype frequencies are reported at antigen recognition domain (ARD) resolution rather than full-field WHO allele resolution. Second, haplotype phasing was inferred using population-based maximum-likelihood methods rather than directly observed through long-read sequencing of the MHC region. Although these methods have been extensively validated and perform well for population-level inference, some uncertainty remains for rare haplotypes, particularly in the presence of residual genotyping ambiguity or incomplete locus coverage at *HLA-DQA1, HLA-DPA1*, and *HLA-DPB1*. As with all EM-based haplotype estimation approaches, uncertainty in rare haplotype inference may influence downstream population genetic diversity metrics and comparative analyses. Finally, population categories were based on self-identified race and ethnicity reported during registry recruitment. While these categories have been shown to correlate strongly with HLA variation and are operationally useful for donor matching and frequency estimation, human genetic variation is continuous and additional substructure likely exists within these groups that cannot be fully captured without genome-wide ancestry informative markers.

Looking ahead, the analytical framework described here can be extended to incorporate additional loci across the MHC region and to integrate emerging high-resolution sequencing data. As long-read sequencing technologies increasingly enable full-gene and phased characterization of HLA haplotypes, future studies will be able to refine these frequency estimates and move toward experimentally resolved MHC haplotypes across populations. Together, these analyses provide the most comprehensive high resolution ARD-level reference of HLA haplotype diversity in US populations to date, establishing a scalable framework for modeling immunogenetic diversity and improving donor–recipient matching in transplantation and related immunogenetic applications.

## ACKNOWLEDGEMENTS

The authors thank the HLA laboratories who contributed genotyping data to the NMDP registry and the over 44 million volunteer stem cell donor registry members worldwide. We would like to recognize the late Captain Robert Hartzman, MD who envisioned this work 40 years ago. We would also like to acknowledge Mike Halagan who did preliminary work on this study. This work was supported by the Office of Naval Research (Grants: N00014-11-1-0339, N00014-12-1-0142, N00014-13-1-0039, N00014-15-1-0848, N00014-16-1-2020, N00014-17-1-2388, N00014-18-1-2045, N00014-19-1-2888, N00014-20-1-2705, N00014-20-1-2832, N00014-21-1-2954, N00014-23-1-2057). LG is supported in part by grants from NIAID R01 AI173095, U01 AI152960, NIDDK R01 DK139241, R01 DK140336, and NHLBI R01 HL179055. MM is supported in part by grants from NIAID R01 AI128775 and R01 AI158861.

## Notes

https://zenodo.org/records/1796993

### Competing Interest Statement

The authors have declared no competing interest.

