## Supplementary Figures for "Classical HLA Allele and Haplotype Frequency Estimates in US Populations"

Supplementary Figure 1. **Clustering analysis**. Clustering analysis using the software CLUTO for the top 100 nine-locus haplotypes across all US populations. The data is row-normalized, and the clusters are based on a neighbor-joining tree of haplotypes according to their profile across the 21 populations. All population acronyms are described in Table 1.

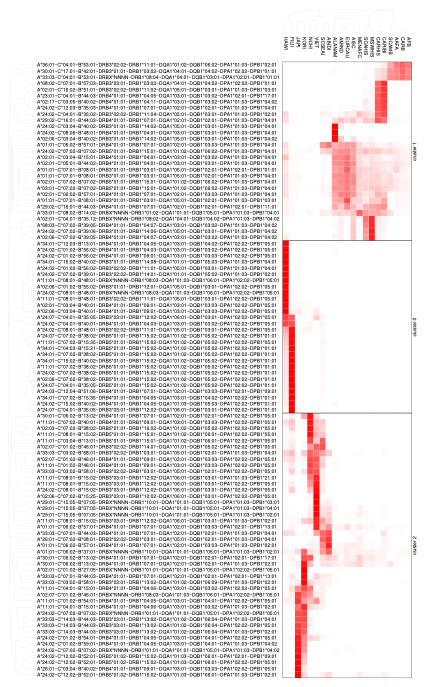

Supplementary Figure 2 - **Population-level HLA allele frequency similarity with the 1000 Genomes Cohort.** Heatmap showing pairwise population similarity between 1000 Genomes (rows) and NMDP (columns) populations based on HLA allele frequency distributions. Similarity is quantified using the Renkonen index, averaged across nine classical HLA loci (HLA-A, -B, -C, -DRB1, -DRB3/DRB4/DRB5, -DQA1, -DQB1, -DPA1, DPB1). Values range from 0 (no overlap) to 1 (identical frequency distributions), with higher values indicating greater similarity. Populations are row-ordered to highlight clusters of maximal correspondence between datasets, illustrating both concordant population structure and differences attributable to sampling, ancestry composition, and registry-specific ascertainment.

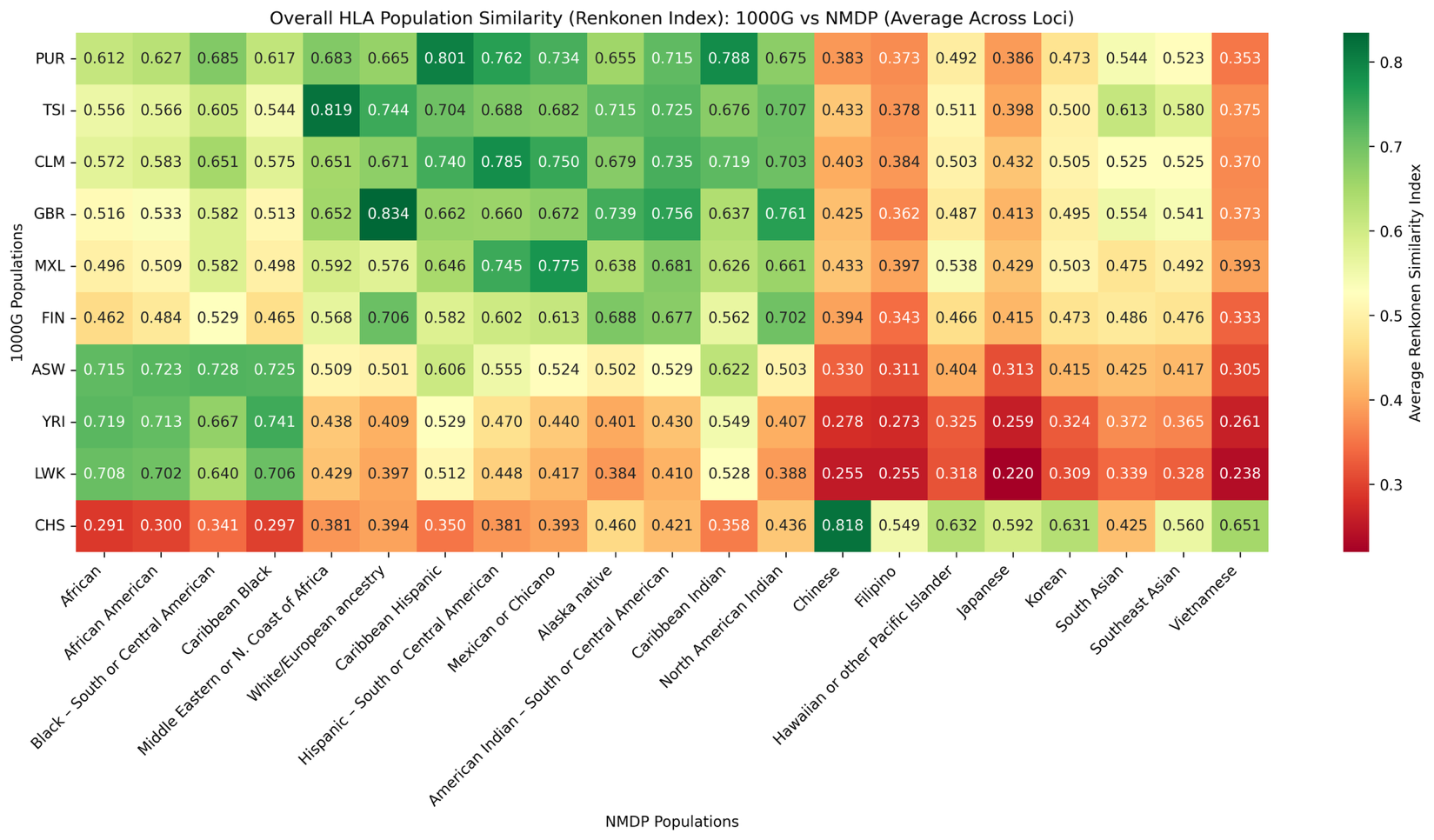

Supplementary Figure 3 - **Haplotype Frequency Comparison for Asian or Pacific Islander broad population category with the Previous 2013 US Haplotype Frequency Publication.** The 40 most common haplotypes are sorted by frequency from the perspective of the current US 2026 dataset with the Y-axis plotted on the log scale.

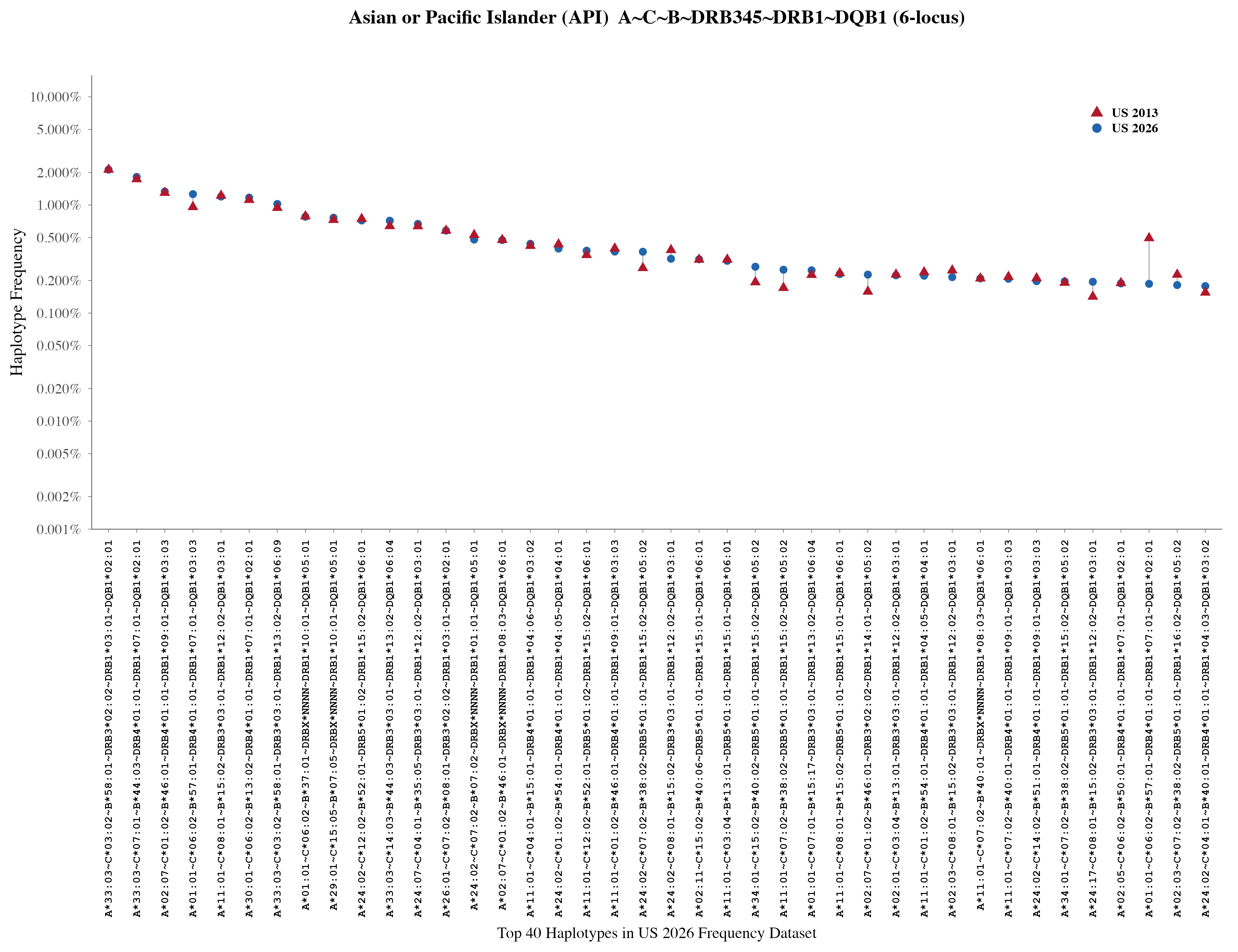
