## Supplementary Methods for "Classical HLA Allele and Haplotype Frequency Estimates in US Populations"

1. **HLA Genotyping Strategies and Reporting Methods:**

Registry HLA typings are updated over time as potential donors are selected for additional typing to confirm high resolution donor-recipient matching. The highest resolution reported typings are used operationally for registry donor selection and for this study. NMDP captures primary data from HLA typing from recruitment laboratories in an XML-based format, Histoimmunogenetics Markup Language (HML) [1]. Primary data captured until 2016 were reinterpreted to IPD-IMGT/HLA database version 3.29.0, after which time HLA genotypes were interpreted to allele list used at time of reporting.

NGS typing and inclusion of the DPB1 locus at recruitment began in 2015, expanding from exon 2 encodes the antigen recognition domain to cover exons 2 and 3 for DRB1/3/4/5, DQB1, and DPB1. Typing of the DQA1 and DPA1 loci at recruitment began in 2020, after which point only exon 2 was sequenced and reported for all class II loci. Before 2015, registry typing methods were more heterogeneous and targeted Sanger sequencing and sequence-specific oligonucleotide (SSO) typing, providing more ambiguous lists of possible HLA genotypes [2].

2. **Limitations of HLA Typing Resolution for Analysis Due to HLA Sequencing Strategy**

The ‘g-group’ designation utilized for HLA allele specificities in this study is not standard WHO nomenclature but was defined in our previous study as a designation combining alleles with identical amino sequence encoded by antigen recognition domain exons (exons 2 and 3 for Class I loci, and exon 2 for Class II loci) [3]. HLA frequency analysis at the “P-group” level was also not achievable, because null alleles could not be systematically excluded by sequencing just exon 2. We also found that historical typings reported to NMDP using P-group designations could lose P-group status after new null alleles were discovered or when there was a change in the current understanding of the expression status of one of the possible HLA alleles.

Three-field “G-group” and full-field WHO allele level analysis was achieved for some but not all registry members, because most reported typing was either stored at two-field level and/or systematic typing of the full gene was not widely performed for HLA Class II loci. The current NMDP donor recruitment HLA typing strategy does not result in consensus sequences that achieve full field WHO allele designation for any HLA locus. For HLA Class I where nearly the full gene is sequenced, reference allele sequences in IPD-IMGT/HLA database extend into portions of the 3’ and 5’ untranslated regions (UTRs) that remain untargeted for sequencing, rendering some four-field allele combinations undistinguished by the typing method.

3. **Managing Copy Number Variation and Limitations of Reporting Methods for HLA-DRB3/4/5 Typings:**

Reporting of typing at DRB3/4/5 loci to NMDP has often not distinguished absence of the gene from lack of typing at a locus. Historic registry typing included an era where neither DRB3 nor DRB4 nor DRB5 were systematically typed and an era where just DRB3 and DRB5 but not DRB4 were typed. We excluded all HLA typings that could not follow standard linkage rules for DRB3/4/5~DRB1 associations [DRB1*03,11,12,13,14 alleles with DRB3, DRB1*04,07,09 alleles with DRB4, DRB1*15,16 alleles with DRB5, and DRB1*01,08,10 with lack of a DRB3/4/5 gene]. Commercial typing kit interpretation software used for registry typing also made such assumptions independently of our process for probe interpretation. When DRB3/4/5 are expected based on the linkage rules, but absent from the reported typing, a genotype list with all possible alleles at that locus was generated and applied to the typing. In the previous study we included heterozygous DRB3/4/5 typings where two genes present but the linkage rules were violated, therefore some exceptions from common linkage are present in the previous dataset, but due to the limitation of reporting we were unable to estimate frequency of absence of DRB3/4/5 gene on haplotypes where it would be expected based on common linkage rules.

A consequence of our analysis strategy to manage this imperfect reporting of DRB3/4/5 typing is that we are unable to capture some Class II haplotypes observed in other datasets that violate the common linkage rules, where a DRB3/4/5 gene is expected but not observed (e.g. DRBX*NNNN~DRB1*15:03~DQB1*06:02, frequently observed in individuals of African ancestry and other common instances where there is an unexpected DRB3/4/5 like a DRB5 allele with DRB1*01:01).

4. **Greedy Algorithm for Per-Population Minimal Allele List Reduction of HLA Typing Data**

High levels of HLA typing ambiguity and continual expansion of alleles in IPD-IMGT/HLA database releases leads to computational challenges for EM haplotype frequency estimation from mixed resolution typing data. Some first-field genotypes with incomplete typing at only A, B, DRB1 may have millions of possible high-resolution genotypes across 9 HLA loci. To address this challenge at the 9-locus resolution, lists of possible alleles were reduced on a per-population basis to only the alleles required for every donor to interpret. We developed and applied a greedy algorithm to determine the minimal list of HLA alleles that could explain every individual in the population. For each locus, we initialized the list with the reference sequence in IPD-IMGT/HLA and attempted to interpret every donor genotype in the population. This minimal allele list generation applied here was different from our previous 2013 [3], where a list of common alleles per population (allele frequency 1/2000 or greater from our 2007 haplotype frequency study [4] were used for initial seeding of the allele list.

Alleles were added to this minimal allele list in an iterative process. For donor genotypes that could not be interpreted using the minimal allele list, we determined which allele should be added to the list next based on counting the number of genotypes where only one additional allele would be needed to interpret the typing. This iterative greedy algorithm continued until all HLA typings could be interpreted for the population.

5. **A Partition-Ligation Approach for Extended 9-Locus Haplotype Frequency Estimation**

The computational challenge of HLA haplotype frequency estimation increases exponentially with the number of loci included. With each additional HLA locus, there is a doubling in the number of possible phased haplotype pairs when given a multi-locus unphased genotype, even when an HLA typing has no allelic ambiguity. Inclusion of HLA typings with allele ambiguity or missing loci poses further challenges, because each possible combination of alleles at a locus must be evaluated.

To address the computational barriers with multi-locus haplotype frequency estimation, we applied a partition-ligation approach where haplotype blocks were iteratively built up using 2-locus EM algorithm. We first estimated haplotype frequencies for a pair of HLA loci, then for each individual, excluded possible haplotype pairs that were not included in 99.9% cumulative probability distribution. Then we treated the haplotype block as a single-locus genotype list and ran a two-locus haplotype frequency estimation to ligate another locus. For HLA Class I we estimated the C~B haplotype first, then combined the A locus to generate A~C~B haplotype frequency estimates. For HLA Class II we started with computing DRB3/4/5~DRB1 two-locus haplotypes then added DQB1, then DQA1. The DPA1~DPB1 block is estimated separately then added to the previous class II DRB3/4/5~DRB1~DQA1~DQB1 aggregate block. Haplotypes for more tightly linked loci with closer genomic distance were combined first, and the more polymorphic beta genes were added first before their pairing with the closely linked alpha genes This strategy was adopted to improve the prediction of the alpha genes due to the increased Asymmetric Linkage Disequilibrium (ALD) in the direction of the Beta loci and the lower coverage of the alpha loci in the study populations. This means knowledge of a beta locus (e.g. DPB1) would lead to better prediction of the alpha locus (e.g. DPA1) than the opposite direction. The Class I haplotype block A~C~B and the DRB3/4/5~DRB1~DQA1~DQB1~DPA1~DPB1 haplotype block were combined in a final ligation step to compute the final 9-locus haplotype frequency estimates.
