## Supplementary Tables for "Classical HLA Allele and Haplotype Frequency Estimates in US Populations"

**Supplementary Table 1. HLA-DQ Group 1 and Group 2 Frequencies Among US Detailed Population Categories.** HLA-DQ Group 1 represents haplotypes containing first-field HLA-DQA1*02/03/04/05/06 alleles. HLA-DQ Group 2 represents haplotypes containing first-field HLA-DQA1*02/03/04/05/06 alleles. Black populations have an equal balance of HLA-DQ Group 1 versus Group 2, while Hispanic populations have a higher frequency of HLA-DQ Group 1 at up to 70.9%.

| **Broad Group** | **Population** | **DQ Group** | **Frequency** |
| --- | --- | --- | --- |
| Black | African American | DQ_G1 | 51.4% |
|  |  | DQ_G2 | 48.6% |
|  | African | DQ_G1 | 50.4% |
|  |  | DQ_G2 | 49.6% |
|  | Black Caribbean | DQ_G1 | 49.6% |
|  |  | DQ_G2 | 50.4% |
|  | Black South or Central American | DQ_G1 | 55.1% |
|  |  | DQ_G2 | 44.9% |
| Asian or Pacific Islander | South Asian | DQ_G1 | 49.4% |
|  |  | DQ_G2 | 50.6% |
|  | Filipino | DQ_G1 | 46.4% |
|  |  | DQ_G2 | 53.6% |
|  | Hawaiian or Pacific Islander | DQ_G1 | 62.2% |
|  |  | DQ_G2 | 37.8% |
|  | Japanese | DQ_G1 | 55.3% |
|  |  | DQ_G2 | 44.7% |
|  | Korean | DQ_G1 | 57.1% |
|  |  | DQ_G2 | 42.9% |
|  | Chinese | DQ_G1 | 63.9% |
|  |  | DQ_G2 | 36.1% |
|  | Southeast Asian | DQ_G1 | 52.2% |
|  |  | DQ_G2 | 47.8% |
|  | Vietnamese | DQ_G1 | 66.4% |
|  |  | DQ_G2 | 33.6% |
| White | White European | DQ_G1 | 59.0% |
|  |  | DQ_G2 | 41.0% |
|  | Middle Eastern or North African | DQ_G1 | 61.6% |
|  |  | DQ_G2 | 38.4% |
| Hispanic | Hispanic Caribbean | DQ_G1 | 62.4% |
|  |  | DQ_G2 | 37.6% |
|  | Mexican | DQ_G1 | 70.9% |
|  |  | DQ_G2 | 29.1% |
|  | Hispanic South or Central American | DQ_G1 | 68.2% |
|  |  | DQ_G2 | 31.8% |
| Native American | Native South or Central American | DQ_G1 | 66.0% |
|  |  | DQ_G2 | 34.0% |
|  | Native Alaska | DQ_G1 | 67.3% |
|  |  | DQ_G2 | 32.7% |
|  | Native North American | DQ_G1 | 67.7% |
|  |  | DQ_G2 | 32.3% |
|  | Native Caribbean | DQ_G1 | 60.8% |
|  |  | DQ_G2 | 39.2% |

**Supplementary Table 2. HLA-DP Amino Acid Level Haplotype Motif Frequencies Among US Detailed Population Categories.**

HLA-DP alpha/beta chain amino acid motif haplotype frequencies and normalized linkage disequilibrium (D') across 21 detailed NMDP populations. The HLA-DP motif is defined by two polymorphic sites: *HLA-DPA1* position 31 (M = methionine or Q = glutamine) in the beta-pleated sheet floor of the peptide-binding groove, and *HLA-DPB1* positions 85–87 in hypervariable region F of the alpha helix. The two major DPB1 motifs are GPM (Gly-Pro-Met) and EAV (Glu-Ala-Val); together with the *HLA-DPA1* position 31 dimorphism, these define four major DPA1~DPB1 haplotype motifs: M~GPM, M~EAV, Q~EAV, and Q~GPM. In a previous NMDP study of the White population, three positions at DPB1 85–87 are in complete linkage disequilibrium and are located in the P1 pocket of the peptide-binding region, influencing peptide-binding specificity and T-cell receptor contact (Hollenbach et al. 2012, Immunogenetics). D' values near +1.0 or −1.0 indicate strong positive or negative LD between the *HLA-DPA1* and *HLA-DPB1* motifs; D' is computed after excluding null and rare ambiguous motifs and renormalizing frequencies. In this study we found while White populations have a very low frequency of the Q~GPM motif and very high D’ values above 0.99, Asian or Pacific Islander populations have a frequency of up to 6.6% for the Q~GPM motif and D’ values as low as 0.7 in the Chinese detailed population.

| **Broad Group** | **Population** | **HLA-DP AA-Level Haplotype Motif** | **Frequency** | **D'** |
| --- | --- | --- | --- | --- |
| Black | African American | M~GPM | 41.1% | 0.998 |
|  |  | M~EAV | 3.0% | -0.998 |
|  |  | Q~EAV | 55.8% | 0.998 |
|  |  | Q~GPM | 0.1% | -0.998 |
|  | African | M~GPM | 44.9% | 0.993 |
|  |  | M~EAV | 4.8% | -0.993 |
|  |  | Q~EAV | 50.2% | 0.993 |
|  |  | Q~GPM | 0.2% | -0.993 |
|  | Black Caribbean | M~GPM | 40.6% | 0.997 |
|  |  | M~EAV | 3.3% | -0.997 |
|  |  | Q~EAV | 56.0% | 0.997 |
|  |  | Q~GPM | 0.1% | -0.997 |
|  | Black South or Central American | M~GPM | 52.9% | 0.999 |
|  |  | M~EAV | 2.3% | -0.999 |
|  |  | Q~EAV | 44.8% | 0.999 |
|  |  | Q~GPM | 0.0% | -0.999 |
| Asian or Pacific Islander | South Asian | M~GPM | 79.8% | 0.893 |
|  |  | M~EAV | 3.3% | -0.893 |
|  |  | Q~EAV | 15.5% | 0.893 |
|  |  | Q~GPM | 1.5% | -0.893 |
|  | Filipino | M~GPM | 10.4% | 0.880 |
|  |  | M~EAV | 2.3% | -0.880 |
|  |  | Q~EAV | 86.1% | 0.880 |
|  |  | Q~GPM | 1.2% | -0.880 |
|  | Hawaiian or Pacific Islander | M~GPM | 44.9% | 0.977 |
|  |  | M~EAV | 2.2% | -0.977 |
|  |  | Q~EAV | 52.4% | 0.977 |
|  |  | Q~GPM | 0.5% | -0.977 |
|  | Japanese | M~GPM | 39.2% | 0.961 |
|  |  | M~EAV | 0.9% | -0.961 |
|  |  | Q~EAV | 58.8% | 0.961 |
|  |  | Q~GPM | 1.1% | -0.961 |
|  | Korean | M~GPM | 41.3% | 0.935 |
|  |  | M~EAV | 1.6% | -0.935 |
|  |  | Q~EAV | 54.2% | 0.935 |
|  |  | Q~GPM | 3.0% | -0.935 |
|  | Chinese | M~GPM | 23.5% | 0.700 |
|  |  | M~EAV | 6.2% | -0.700 |
|  |  | Q~EAV | 63.7% | 0.700 |
|  |  | Q~GPM | 6.6% | -0.700 |
|  | Southeast Asian | M~GPM | 62.4% | 0.911 |
|  |  | M~EAV | 4.9% | -0.911 |
|  |  | Q~EAV | 30.8% | 0.911 |
|  |  | Q~GPM | 1.9% | -0.911 |
|  | Vietnamese | M~GPM | 22.0% | 0.683 |
|  |  | M~EAV | 12.4% | -0.683 |
|  |  | Q~EAV | 59.8% | 0.683 |
|  |  | Q~GPM | 5.8% | -0.683 |
| White | White European | M~GPM | 76.4% | 0.989 |
|  |  | M~EAV | 11.9% | -0.989 |
|  |  | Q~EAV | 11.6% | 0.989 |
|  |  | Q~GPM | 0.1% | -0.989 |
|  | Middle Eastern or North African | M~GPM | 80.2% | 0.944 |
|  |  | M~EAV | 8.5% | -0.944 |
|  |  | Q~EAV | 10.8% | 0.944 |
|  |  | Q~GPM | 0.5% | -0.944 |
| Hispanic | Hispanic Caribbean | M~GPM | 81.2% | 0.998 |
|  |  | M~EAV | 5.1% | -0.998 |
|  |  | Q~EAV | 13.6% | 0.998 |
|  |  | Q~GPM | 0.0% | -0.998 |
|  | Mexican | M~GPM | 84.3% | 0.995 |
|  |  | M~EAV | 6.1% | -0.995 |
|  |  | Q~EAV | 9.5% | 0.995 |
|  |  | Q~GPM | 0.0% | -0.995 |
|  | Hispanic South or Central American | M~GPM | 82.8% | 0.991 |
|  |  | M~EAV | 5.2% | -0.991 |
|  |  | Q~EAV | 11.9% | 0.991 |
|  |  | Q~GPM | 0.1% | -0.991 |
| Native American | Native South or Central American | M~GPM | 82.8% | 0.983 |
|  |  | M~EAV | 6.2% | -0.983 |
|  |  | Q~EAV | 10.8% | 0.983 |
|  |  | Q~GPM | 0.2% | -0.983 |
|  | Native Alaska | M~GPM | 76.3% | 0.966 |
|  |  | M~EAV | 10.4% | -0.966 |
|  |  | Q~EAV | 13.0% | 0.966 |
|  |  | Q~GPM | 0.3% | -0.966 |
|  | Native North American | M~GPM | 86.2% | 0.996 |
|  |  | M~EAV | 6.5% | -0.996 |
|  |  | Q~EAV | 7.3% | 0.996 |
|  |  | Q~GPM | 0.0% | -0.996 |
|  | Native Caribbean | M~GPM | 69.8% | 0.993 |
|  |  | M~EAV | 7.9% | -0.993 |
|  |  | Q~EAV | 22.2% | 0.993 |
|  |  | Q~GPM | 0.1% | -0.993 |

**Supplementary Table 3. Haplotype Frequency Concordance: US2026 vs. US2013**

Six-locus (A~C~B~DRB345~DRB1~DQB1) haplotype frequency comparison between the current US2026 dataset and the Gragert et al. 2013 published dataset. The NMDP donors included in US2013 overlap with those in the US2026 dataset.

| **Population** | **N (US2026)** | **N (Prev)** | **If (%)** | **If SE (%)** |
| --- | --- | --- | --- | --- |
| Black | 938,006 | 505,223 | 83.8% | 92.9% |
| Asian or Pacific Islander | 1,081,437 | 568,579 | 82.3% | 90.7% |
| White | 6,247,346 | 3,912,408 | 88.0% | 91.5% |
| Hispanic | 1,506,537 | 712,736 | 83.5% | 91.1% |
| Native American | 115,550 | 46,137 | 63.7% | 81.8% |
| African American | 696,800 | 416,560 | 83.6% | 93.0% |
| African | 67,127 | 28,550 | 63.3% | 89.4% |
| Black Caribbean | 62,075 | 33,320 | 71.0% | 93.0% |
| Black South or Central American | 7,672 | 4,885 | 49.2% | 89.9% |
| South Asian | 349,054 | 185,382 | 82.9% | 92.7% |
| Filipino | 94,813 | 50,608 | 81.2% | 92.6% |
| Hawaiian or Pacific Islander | 21,351 | 11,498 | 69.4% | 89.3% |
| Japanese | 36,700 | 24,581 | 79.8% | 93.3% |
| Korean | 127,213 | 77,582 | 82.5% | 92.5% |
| Chinese | 194,210 | 99,666 | 80.8% | 92.1% |
| Southeast Asian | 57,668 | 27,971 | 66.9% | 89.5% |
| Vietnamese | 84,799 | 43,538 | 81.6% | 93.3% |
| White European | 3,118,526 | 1,242,874 | 90.1% | 94.4% |
| Middle Eastern or North African | 182,812 | 70,880 | 75.5% | 92.6% |
| Hispanic Caribbean | 172,424 | 115,366 | 80.8% | 92.3% |
| Mexican | 463,491 | 261,218 | 82.8% | 92.4% |
| Hispanic South or Central American | 261,463 | 146,704 | 79.6% | 92.9% |
| Native South or Central American | 10,329 | 5,921 | 60.1% | 91.7% |
| Native Alaska | 5,026 | 1,374 | 43.5% | 86.0% |
| Native North American | 52,011 | 35,781 | 77.3% | 92.7% |

**Supplementary Table 4. Haplotype Frequency Concordance with Published NMDP HLA Class II Datasets**

Renkonen If metric comparing US2026 haplotype frequencies with previously published NMDP US high-resolution typed datasets incorporating HLA Class II DQA1 or DPB1 loci. If represents the proportion of the frequency distribution shared between two datasets (1.0 = identical). If SE is the lower-bound If when accounting for standard error of individual haplotype frequency estimates. The NMDP donors included in these published datasets overlap with those in the US2026 dataset.

| **Comparison** | **HLA Loci** | **Population** | **N (US2026)** | **N (Prev)** | **If (%)** | **If SE (%)** |
| --- | --- | --- | --- | --- | --- | --- |
| US2026 vs. Klitz et al. 2003 | DRB1~DQA1~DQB1 | White | 6,247,346 | 1,899 | 91.7% | 95.5% |
| US2026 vs. Hollenbach et al. 2012 | DPA1~DPB1 | White | 6,247,346 | 5,944 | 89.7% | 91.6% |

**Supplementary Table 5. Haplotype Frequency Concordance with IHIW Datasets**

Comparison of 6-locus (A~C~B~DRBX~DRB1~DQB1) and 9-locus (A~C~B~DRBX~DRB1~DQA1~DQB1~DPA1~DPB1) haplotype frequencies with 17th International HLA and Immunogenetics Workshop datasets (<https://17ihiw.org/17th-ihiw-ngs-hla-data/>) and Creary et al. 2019 for the White population. “Freq Sum” indicates the fraction of the total frequency distribution present in each dataset, as IHIW datasets do not report the haplotypes where there is higher uncertainty (count <3), the “Freq Sum IHIW” values are substantially less than 100% after trimming.

| **Comparison** | **HLA Loci** | **Population** | **N (US2026)** | **N (IHIW)** | **If (%)** | **If SE (%)** | **Freq Sum US2026 (%)** | **Freq Sum IHIW (%)** |
| --- | --- | --- | --- | --- | --- | --- | --- | --- |
| US2026 vs. Creary et al. 2019 | 9-locus | White | 6,247,346 | 633 | 56.9% | 65.1% | 100% | 31% |
| US2026 vs. 17th IHIW | 6-locus | Asian or Pacific Islander | 1,081,437 | 12 | 36.1% | 54.4% | 100% | 31% |
| US2026 vs. 17th IHIW | 6-locus | White | 6,247,346 | 1,490 | 62.7% | 74.7% | 100% | 68% |
| US2026 vs. 17th IHIW | 6-locus | Black | 938,006 | 131 | 44.3% | 61.3% | 100% | 35% |
| US2026 vs. 17th IHIW | 6-locus | Hispanic | 1,506,537 | 13 | 37.3% | 54.8% | 100% | 27% |
| US2026 vs. 17th IHIW | 9-locus | Asian or Pacific Islander | 1,081,437 | 11 | 36.7% | 58.4% | 100% | 28% |
| US2026 vs. 17th IHIW | 9-locus | White | 6,247,346 | 1,047 | 56.0% | 68.7% | 100% | 48% |
| US2026 vs. 17th IHIW | 9-locus | Black | 938,006 | 60 | 46.1% | 62.1% | 100% | 16% |
| US2026 vs. 17th IHIW | 9-locus | Hispanic | 1,506,537 | 9 | 41.4% | 57.6% | 100% | 19% |
